# Compartmentalized above- and belowground defenses in *Tanacetum vulgare* are tailored to localized antagonists

**DOI:** 10.1101/2025.10.30.685307

**Authors:** Humay Newrzella, Robin Heinen, Cecilia M. Villasante, Ina Zimmer, Baris Weber, Peter Kary, Georg Gerl, Annika Neuhaus, Alexandros Sigalas, Lina Ojeda-Prieto, Jana Barbro Winkler, Wolfgang W. Weisser, Jörg-Peter Schnitzler

## Abstract

Specialized metabolites, specially terpenoids, play a key role in plant defense. However, how terpenoid diversity governs inducible chemistry and root architectural development remain poorly understood. We used a combination of high-throughput root phenotyping and targeted metabolite profiling to examine three leaf terpenoid chemotypes of common tansy (*Tanacetum vulgare*). Using a phenotyping platform, we tested whether (i) root-chewing wireworms induce root terpenoids locally and alter shoot terpenoids systemically, (ii) phloem-feeding aphids elicit chemotype-dependent responses, and (iii) chemotypes differ in root-system development. After root establishment, the plants were exposed to wireworms (*Agriotes* spp.) and aphids (*Macrosiphoniella tanacetaria*), both separately and together, and were then monitored for 60 days. The chemotypes differed in inducible chemistry and root architecture. Chemotype 1 developed the fastest-growing root systems and the highest root:shoot ratios. Wireworms increased stored root sesquiterpenoid levels by more than twofold in chemotypes 1 and 2, whereas chemotype 3 was largely unresponsive. Aphids didn‘t alter root terpenoids, but significantly increased leaf monoterpenoid emissions in chemotype 1 without affecting stored pools. Therefore, storage and emission were decoupled and depended on both organ and chemotype. Our analysis reveals a compartmentalized, chemotype-specific defense strategy in tansy, highlighting the coordinated regulation of the root system and inducible chemistry.

**Highlight:** In *Tanacetum vulgare*, wireworms boost root sesquiterpenoids and aphids elevate leaf monoterpenoid emissions; chemotype governs terpenoid defense and root system architecture.

## Introduction

The plant kingdom produces a massive range of specialized metabolites that convey chemical information among plants, insects, microorganisms, and adjacent roots (Massalha et al., 2017; Erb & Reymond, 2019). These metabolites function in defense, prime uninjured tissues and neighboring plants, regulate rhizosphere microbial populations, and affect plant fitness across ecological levels (Heil & Karban, 2010; Frost et al., 2008; Kong et al., 2021).

Terpenoids, specifically monoterpenoids (C₁₀) and sesquiterpenoids (C₁₅), constitute a major fraction of this chemical diversity in plants (Gershenzon & Dudareva, 2007; Tholl, 2015). Mono- and sesquiterpenoids are synthesized from geranyl- or farnesyl-diphosphate via the enzyme activity of terpenoid synthases (TPSs), which can convert a single substrate into multiple products, thereby substantially expanding structural diversity in plants (Tholl, 2015; Christianson, 2017). Upon their formation, terpenoids serve multiple roles in defense, and many of them exhibit direct toxic or deterrent effects against herbivores and pathogens (Gershenzon & Dudareva, 2007). Herbivore-induced terpenoid blends emitted from damaged foliage may also provide an indirect line of defense by attracting predators and parasitoids to the affected plants (Aartsma et al., 2017; Turlings et al., 1990). A similar concept also applies to the belowground environment. For instance, maize roots under larval attack release the sesquiterpenoid (E)-β-caryophyllene, which in turn attracts entomopathogenic nematodes that infest the larvae, and thereby mitigates damage from root-feeding beetles (Rasmann et al., 2005, Degenhardt et al., 2009). Terpenoids are induced in tissues via two mechanisms. Either they are actively emitted from plants by abiotic and biotic stress or they can also be released following disruption of reservoir glands (Niinemets et al., 2013; Vickers et al., 2009). In general, terpenoids are stored in reservoir glands found in most plant species (Lange and Turner, 2013). For instance, a species like *Tanacetum vulgare* possesses glandular trichomes in which they accumulate terpenoids in leaf tissues (Guerreiro et al., 2016). Likewise, many *Lamiaceae* (mint family) species have abundant peltate glandular trichomes whose essential oils are largely terpenoids (Schuurink and Tissier, 2020). In contrast, precise storage sites of terpenoids in roots are still poorly resolved for many species. Among the few known storage locations are the lactifers in the roots of dicots such as dandelion that sequester defensive sesquiterpene lactones (Huber et al., 2016). Terpenoids stored in leaf tissues exhibit significant intraspecific variation and can be used for classifying plant chemotypes. Chemotypes are typically characterized by a single dominant compound or by multiple combinations that do not exhibit distinct dominant substances, resulting in mixed chemotypes (Clancy et al., 2016; Neuhaus-Harr et al., 2024; Rahimova et al., 2024). Chemotype composition has also been connected to various ecological functions in plant-insect interactions, including defense strategies. For instance, leaves in tea-tree (*Melaleuca alternifolia*) plants are classified into terpenoid chemotypes, distinguished by the predominance of cineole, terpinolene, and terpinen-4-ol (Bustos-Segura et al. 2015). The same study additionally revealed that *Faex sp*. larvae grew faster on cineole-rich plants, *P. tigrina* adults damaged terpinolene-rich plants less, and cineole-rich plants were more susceptible to myrtle rust *(Puccinia psidii).* These results show that within species terpenoid chemotypes might shape plant-herbivore and plant-pathogen interactions aboveground. By contrast, far less is known about terpenoid storage sites and its dynamics and roles in roots, partly because belowground volatiles are hard to sample and quantify in soil, and microbial turnover can confound measurements (Delory et al., 2016). Nevertheless, a few studies demonstrate ecological functions of root terpenoids. For instance, in cotton *(Gossypium herbaceum),* terpenoid aldehydes are induced in roots after herbivory damage (Bezemer et al., 2004); in common dandelion *(Taraxacum officinale)*, a sesquiterpene lactone in root latex deters cockchafer larvae *(Melolontha melolontha)* and improves plant fitness (Huber et al., 2016); and in *Centaurea stoebe,* root sesquiterpene emissions modify neighbors and can increase root herbivore performance on adjacent plants (Gfeller et al., 2019; Huang et al., 2019). These examples suggest that root terpenoids can be ecologically consequential, yet comprehensive datasets linking genotype-specific root terpenoid profiles to herbivore impacts remain scarce. At a mechanistic level, many root terpenoids are jasmonic acid (JA)-dependent (Tholl, 2015), whereas phloem-feeding aphids typically do not activate JA signaling in roots (Karssemeijer et al., 2020). Accordingly, foliar aphid attack is not expected to elicit a systemic increase in root sesquiterpenoids. By contrast, aphids can cause modest, chemotype-dependent rises in foliar monoterpene emissions (Clancy et al., 2020).

Chemical diversity in plants defines not only chemotype profiles but can also be associated with growth traits. For instance, Ojeda-Prieto et al. (2024) documented that in common tansy *(Tanacetum vulgare)*, foliar terpenoid chemotypes were associated with early differences in growth that converged over the season, whereas chemotype effects on reproduction (flower-head number and flowering phenology) persisted, linking terpenoid chemistry to reproductive performance. Comparably, in *Cinnamomum camphora*, predefined leaf-oil chemotypes (linalool, eucalyptol, camphor, and borneol) differed in leaf morphology and photosynthetic performance. Linalool and eucalyptol chemotypes had larger leaves, and eucalyptol and camphor chemotypes showed higher photosynthetic rates (associational differences among chemotype groups, Luo et al., 2021). Root metabolomic composition has also been associated with root architecture in crops (Ghorbanzadeh et al., 2023). Because comparable datasets for terpenoid-rich species remain scarce, we test whether root system architecture varies among chemotypes.

To study inducible chemistry in above- and belowground systems and root development in terpenoid-rich species, we used tansy *(Tanacetum vulgare)*, a species whose individuals differ strikingly in terpenoid composition and can therefore be assigned to distinct chemotypes. These chemotypes have so far been defined mainly by monoterpenoids, which are abundant in leaves and are known to influence interaction between tansy and associated insects (Clancy et al., 2016; Neuhaus-Harr et al., 2024; Neuhaus-Harr et al., 2025). Our recent work shows, however, that sesquiterpenoids are just as integral to tansy chemistry as monoterpenoids with potential different ecological functions (Rahimova et al., 2024). For example, earlier we demonstrated sharp organ-specific sesquiterpenoid profiles across leaves, rhizomes, coarse roots, and fine roots in tansy (Rahimova et al., 2025). Furthermore, root application of pipecolic acid (a systemic acquired resistance cue) strongly induced multiple root sesquiterpenoids, significantly more than in shoots. Yet whether root sesquiterpenoid chemotypes are similarly inducible under real herbivory remains unknown. To fill that gap, we exposed three chemotypes to both herbivores in sequence – root-chewing wireworms *(Agriotes spp.)* followed by the phloem-feeding aphid *(Macrosiphoniella tanacetaria)* – and quantified terpenoid stores and emissions from leaves and roots to test how cross-guild attacks influence chemotype-dependent defenses above- and belowground. We asked the following questions:

**1.** Do both wireworms and aphids, applied sequentially, trigger localized vs. systemic shifts in terpenoid stores in tansy roots and shoots, and are the magnitudes of these shifts chemotype-specific?
**2.** Does feeding by the aphids, either alone or together with wireworms, change terpenoid emission in shoots, and are these changes chemotype-specific?
**3.** Do plant phenotype traits, specifically root system architecture, differ depending on the chemotype identity in tansy.

## Material and Methods

### Plant material

Three *Tanacetum vulgare* chemotypes were selected from a stock of chemotypes available at the Terrestrial Ecology Research Group, and originally sourced from seeds collected in Jena, Germany. For further details of the origin of the chemotypes and thus plant materials, please see Neuhaus-Harr et al., 2024. All plants were propagated vegetatively via stem-cuttings taken from plants in the end of summer prior to the experiment which took place in autumn. The cuttings were potted in soil and grown for two weeks in controlled conditions with a photoperiod of 14 h:10 h and a temperature of 21°C:17°C (day:night).

### Experimental set-up

In order to assess the phenotypic development of tansy and the effect of above- and belowground treatments in a non-invasive approach, we used rhizotrons in a high-throughput phenotyping (HTP) facility (Fitness-SCREEN) at Helmholtz Munich. Two-week-old plants from three distinct chemotypes were transferred to the designated rhizotrons (40 cm × 80 cm × 4 cm; length × height × width) containing a peat substrate (Einheitserde Classic type CL-T, Patzer Erden GmbH, Germany) and black basalt sand mixture (1/1.2, v/v). The plants were cultivated under controlled conditions, with a 14-hour photoperiod (light:dark) at 20°C:18°C and 40%:60% relative humidity (day:night). Prior to the treatment, each plant was watered twice a week with 200 mL of the diluted (1/100, v/v) fertilizer Hakaphos® Rot (8% N, 12% P2O5, 24% K2O, 4% MgO, 31% SO3, 0.01% B, 0.02% Cu, 0.05% Fe, 0.05% Mn, 0.001% Mo, 0.02% Zn; Compo Expert GmbH, Germany). Overall, we transplanted 60 plants (one per rhizotron) into individual rhizotrons positioned on automated carriers. Chemotype identity was randomized across carriers.

### Herbivory and insect colonies

The aboveground treatment involved the use of aphids as phloem-feeding herbivores. The aphid species, *Macrosiphoniella tanacetaria* were obtained from the Terrestrial Ecology Research Group at TUM, and originally collected from plants in Jena, Germany. The aphids were reared on tansy plants, used in our previous study, under controlled environmental conditions (photoperiod 14 h:10 h; 21°C:17°C; day:night). For the plant treatment, age-specific cohorts of aphids were established to maintain comparable population growth across the different plants during the experiment. For this, fifty adult aphids were selected from the main colony and kept on clipped tansy leaves in a petri dish (150 mm in diameter and 15 mm in height) for 24 hours. After removing the adult aphids, the nymphs were allowed to grow for an additional two days, and these two days old nymphs were used for the aphid treatment.

For the belowground treatment, wireworms, which are known to feed on plant roots, were used. The larval population were collected in the fields, and hence consisted of a mixture of *Agriotes lineatus* and *Agriotes obscurus* obtained from Wageningen University in the Netherlands. These larvae were reared in covered soil containers with commercially available potatoes until treatment.

### Treatment

We initiated the treatment 7 weeks after propagation of the plants. The experimental design of our treatment process is provided in Supplementary Figure S1. The treatment groups included control plants with no treatment, plants treated only with wireworms, plants treated only with aphids, and plants that had been treated with both wireworms and aphids. The selection and introduction of worms was conducted in two steps. In the first step, we carefully opened the rhizotrons and randomly placed five worms within the rhizotrons. In the second step, we gently dug small holes in the soil closer to the stem and introduced the next batch of four worms (in total, nine worms per plant) to the soil. We continuously monitored each plant to ensure that worms did not escape and were contained within the rhizotrons.

Ten days after worm treatment, 10 aphid nymphs were carefully applied to one mature leaf of the plants designated for aphid and worm & aphid treatments, and covered with sheer organza bags (9.5 × 15 cm, length × width) to prevent potential escape. After 19 days of the aphid treatment, root and leaf tissue were harvested and frozen in liquid nitrogen. Samples were kept at -70 °C until further chemical analyses. During the final harvest, the number of aphids per leaf was counted and the wireworms were retrieved, except for two that were found dead. At the final harvest, we also assessed the natural infestation level by thrips, powdery mildew, and *Coloradoa tanacetina* that occurred in the greenhouse during the experiment, so that their impacts could be accounted for in the statistical analyses.

### Hexane extraction of terpenoids and GC-MS analyses

We analyzed leaves from 59 plants (chemotype 1: n (biological replicates) = 20, chemotype 2: n = 19, and chemotype 3: n = 20), and roots from 58 plants (chemotype 1: n = 19, chemotype 2: n = 20, and chemotype 3: n = 19). In the laboratory, the frozen leaf and root samples were ground into powder and stored at -70°C until extraction. We conducted the extraction in the following steps: 300 µl hexane + 860 pmoL·μl^−1^ internal standard (monoterpenoid δ-2-carene) was added to approximately 150 mg frozen tissue (leaf or root) powder, the sample was then vortexed for a minute and then stored at 4°C for 24 hours. After 24 hours, 150 μl of the liquid extract was removed and stored at 4°C. Additional 150 μl hexane and internal standard mixture was added to the material, which was then vortexed and stored again at 4°C for 24 hours. 150 μl of the liquid extract was again removed and combined with the previously collected extract. One μl of sample was injected into an empty glass cartridge containing a glass micro-vial that was placed on the autosampler of the gas chromatography mass-spectrometer (GC-MS) system. The GC-MS temperature program, quantification and qualification methods are the same as described in our previous studies (i.e., Rahimova et al., 2024, 2025).

### VOC Collection and GC-MS analyses

During the experiment, VOCs were collected from the root and shoot tissues of plants in the greenhouse using a passive sampling method with Twisters (Gerstel, Mülheim an der Ruhr, Germany) that are made of non-polar polydimethylsiloxane-coated stir bars (film thickness 0.5 mm, length 10 mm). Their non-polar nature allows them to efficiently capture the VOCs. For the VOCs collection from shoot tissues, we attached each sampling twister to paper clips and carefully placed them in the bushy parts of the plants for a duration of 36 hours. Additionally, we positioned twisters in various appropriate locations throughout the greenhouse to ensure a comprehensive background measurement, allowing us to correct later for any potential peaks that might not originate from tansy. For aboveground passive VOC sampling, we analyzed 58 plants (chemotype 1: n=18; chemotype 2: n=20; chemotype 3: n=20). When collecting VOCs from root tissues, we carefully opened the rhizotrons glass covers and inserted twisters into four different sections of the rhizotron and root system. The first twister was attached to the upper part of the rhizotron (approximately 5 cm from the top), where most of the coarse roots are found. The second and third twisters were attached to individual roots, which were gently wrapped in aluminum bags (8 x 5 cm length × width) along with the twisters. These two twisters were positioned in the lower part of the rhizotron, at a distance of 30-40 cm from both the upper and lower part of the rhizotrons. For each rhizotron, a spoonful of soil was also collected in an aluminum bag together with a twister to control for potential contamination from the soil-derived VOCs. The root volatile collection was conducted for a duration of 60 hours.

The headspace analysis of VOC samples was conducted following methodology described by Zhu et al. (2025) using thermo-desorption gas chromatography-mass spectrometry (TD-GC-MS; TD by Gerstel; GC: 7890A, MS: 5975C, Agilent Technologies, Palo Alto, CA, USA). For quantitative and qualitative analyses of the compounds from both methods, solvent extraction and headspace analysis, we used Agilent GC-MS Enhanced ChemStation, version E.02.00.493 (Agilent, St. Clara, USA). The peaks were initially assessed for chromatogram quality based on peak purity. The detected peaks were subsequently identified by comparing the mass spectra of each chromatogram with the Mass Spectral Library (National Institute of Standards and Technology: NIST 20) and the Kovats retention indices. The latter were calculated based on retention times of a saturated alkane mixture (C9-C21; Sigma-Aldrich) measured along with the samples. The quantification of the compounds was carried out using six dilutions of pure standards of sabinene, α-pinene, linalool, methylsalicylate, bornyl acetate, β-caryophyllene, and α-humulene with the concentration range of 55, 120, 250, 400, 550, and 800 pmol/µl.

### Phenotypical analysis

We used a non-invasive, high-throughput phenotyping (HTP) workflow to quantify root traits over 60 days of monitoring. First, image acquisition was performed for belowground traits. The transparent side of each rhizotron was photographed daily with fixed camera setups: Allied Vision Prosilica GT 6600 paired with a Milvus 2/50 M lens, producing 6012 × 2821-pixel grayscale images that spanned the full 685 × 322 mm observation window (≈8765 pixels mm⁻² resolution). Root image analysis pipeline is as follows: first, raw root images (.bmp) were converted to .tif to be compatible with RootPainter 0.2.21 (Smith et al. 2022), an AI-driven segmentation tool that relies on convolutional neural networks (CNNs). Following the corrective training protocol proposed by Seethepalli et al. (2021), RootPainter automatically extracted one 900-pixel tile per image. The operator iteratively annotated roots and background until model predictions stabilized, at which point training was halted and the best-performing model was used to segment the full image set. Segmented outputs were then converted into a format readable by RhizoVision Explorer 2.0.3 for quantitative trait extraction (Seethepalli & York, 2020; Seethepalli et al., 2021).

Within RhizoVision, the “broken roots” mode was selected to improve reconstruction of discontinuous root segments, and the following global filters were applied: threshold = 200, removal of non-root objects larger than 1 mm², and root-pruning threshold = 1. Batch processing produced many features for every image and three of those features, total visible root length, root volume, and root surface area, are used in this study. Shoot architectural features will not be part of this study.

### Statistical analysis

Data visualization was performed in R (version 4.3.3; R Core Team 2024) using ggplot2 (Wickham, 2016) for graphics; patchwork (Pedersen, 2020) to combine panels (e.g. boxplots); corrplot (Wei & Simko, 2017) for correlation matrices; ComplexHeatmap (Gu et al., 2016) for heatmap. Final figures were edited and exported at high resolution in Inkscape, an open-source vector-graphics editor.

#### Analysis of the stored and emitted compounds

To effectively examine the primary focus of our study, we analyzed total concentration of terpenoids (both mono- and sesquiterpenoids) in roots and shoots (stored pools) and shoot emitted terpenoids with linear models in R (base *lm*). Response variables were log-transformed to stabilize variance. Unless noted otherwise, models included the unplanned biotic covariates thrips, powdery mildew, and *Coloradoa tanacetina* (monitoring recorded at final harvest), and the main effect of experimental factors, wireworms (Worm), aphids (Aphid), and plant chemotype, with all two- and three-way interactions among Worm, Aphid and Chemotype. Because stored root monoterpenoids were detected in only 28 out of 58 plants, we excluded zero observations prior to modeling and this created empty cells for some chemotype × treatment cells, yielding a rank-deficient design (aliased, non-estimable coefficients) for terms involving chemotype. We therefore fit a reduced model excluding chemotype and its interaction terms for stored root monoterpenoids. Similarly, emitted sesquiterpenoids were detected in 35 plants out of 58, and thus we modelled the positive subset only. For emission data, the unplanned covariates were omitted because headspace sampling occurred 10 days before harvest (timing mismatch with the covariate measurements). Stored root sesquiterpenoids, stored shoot mono and sesquiterpenoids, and emitted monoterpenoids were positive across all plants and required no special handling. We used Type III ANOVA *(car::Anova;* Fox & Weisberg, 2019) with sum-to-zero contrasts for factors to test model terms. Model assumptions (normality and homoscedasticity of residuals) were assessed by visual inspection of Q-Q and residual-vs-fitted plots.

For the figures presented in this study, we also obtained pairwise comparisons from the full model using estimated marginal means *(emmeans;* Lenth, 2024). We report: 1) all-pairs treatment comparisons or contrasts among the four treatment combinations within a chemotype for Figure 1 (e.g., Control, Worm, Aphid, Worm & Aphid) using Tukey-adjustment; 2) simple-effects within chemotype for Figure 4 (e.g., Aphid vs Control within each chemotype) using *emmeans*, averaging over Worm with observed cell weights *(weights = “cells”)*. P-values were Bonferroni-adjusted across the set of chemotype-specific tests. Statistical significance was set at α = 0.05 after adjustment.

**Figure 1:**
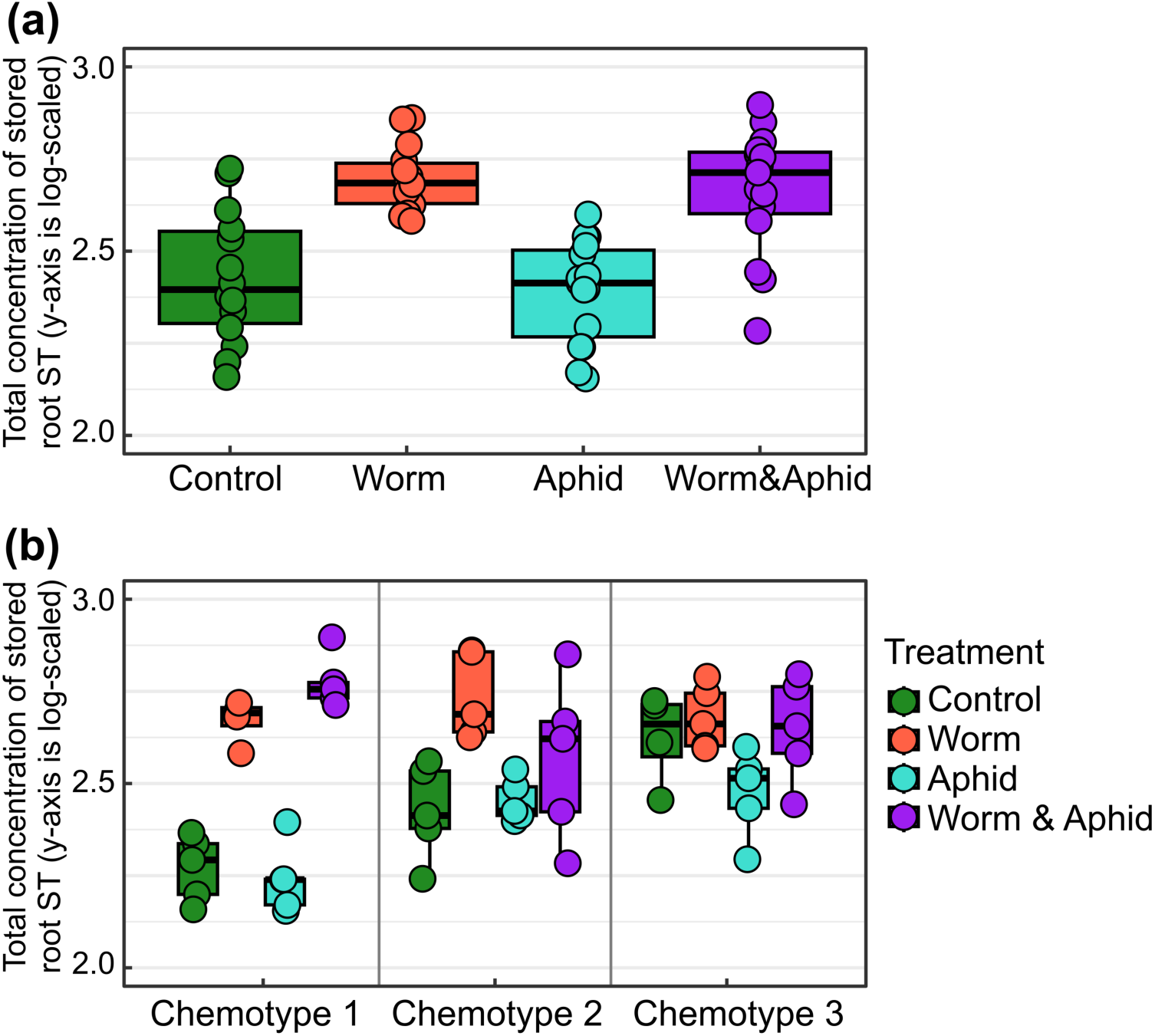
**(a)** Total concentration of stored root sesquiterpenoids (ST) under Control (n = 14), Worm (n = 14), Aphid (n = 15), and Worm&Aphid (n = 15) treatments. Statistical contrasts among treatments: Worm > Control (p < 0.001), Worm&Aphid > Control (p < 0.001), Worm > Aphid (p < 0.001), Worm&Aphid > Aphid (p < 0.001); Aphid vs Control (p = 0.754) and Worm vs Worm&Aphid (p = 0.688) were not significant. **(b)** Chemotypic effect of stored root STs to above and belowground feeding herbivories. Within-chemotype contrasts: Chemotype 1 – Worm > Control (p < 0.001), Worm&Aphid > Control (p < 0.001), Worm > Aphid (p < 0.001), Worm&Aphid > Aphid (p < 0.001); Chemotype 2 – Worm > Control (p < 0.001) and Worm > Aphid (p = 0.001), other pairs not significant; Chemotype 3 – Worm > Aphid (p = 0.014), other pairs not significant. All pairwise contrasts were obtained from the full model via *emmeans* (Tukey-adjustment for multiple comparisons). For panel (a) contrasts are averaged over chemotypes, and panel (b) contrasts are computed within each chemotype. The y-axis is on a logarithmic scale. The boxes enclose the middle half of the data, with the lower edge at the 25th percentile and upper edge at the 75th percentile. The horizontal line inside each box is the median. The whiskers stretch to the most distant points within 1.5 × the box’s height (the interquartile range).

#### Analysis of root-growth traits

We analyzed surface area, total root length, volume, and root:shoot ratio with generalized additive mixed models (GAMMs) fit with *“mgcv”* package (Wood, 2017) and smoothing parameters were estimated by restricted maximum likelihood (REML). To account for repeated measurements of the same plant, we added a plant-level random intercept. For chemotype differences, we obtained estimated marginal means (EMMs) from the GAMMs using *“emmeans”*, averaging uniformly over the observed day-of-year range. Pairwise chemotype contrasts were tested with Wald χ² (df = 1) derived from the EMM contrasts, and p-values were adjusted within trait using the Benjamini-Hochberg (BH) procedure (FDR α = 0.05). We visualized trait trajectories over day of year using *“geom_smooth”* method with *“mgcv::gam”* (thin-plate splines, k = 10, REML), fitting a separate smoother for each chemotype and displaying 95% confidence intervals.

## Results

The monoterpenoid composition of each of the selected three chemotypes was highly distinct (Figure S2a). Leaf tissues of chemotype 1 was β-thujone-dominant, chemotype 2 exhibited a mixed profile with camphor, sabinene hydrate, bornyl acetate, and other minor constituents, and chemotype 3 was dominated by trans-chrysanthenyl acetate with cis-chrysanthenol as a secondary component. These patterns are in alignment with the identities reported for our source material of chemotypes (Athu_Bthu, Mixed_low, Chyrs_Acet) by Neuhaus-Harr et al. (2024, 2025). We note minor quantitative differences, most notably in chemotype 1 where β-thujone accounted for 88% of total monoterpenoids and α-thujone for 3%, whereas Neuhaus-Harr et al. reported a higher combined α-/β-thujone contribution. Such shifts are plausibly attributable to differences in GC-MS methods and extraction protocols between the studies. Importantly, the qualitative profile of each chemotype remained unchanged, supporting the same chemotype assignment. The sesquiterpenoids content in leaf tissues of each chemotype was majorly characterized by the abundance of β-copaene (Figure S2b). Each chemotype, however, also showed a unique sesquiterpenoid, with sabinol isovalerate identified in chemotype 1, β-selinenol in chemotype 2, and spirojatamol in chemotype 3 (Figure S2b). Across the stored root terpenoid profile, even though monoterpenoids were scarce (four compounds detected), their abundance was specific to each chemotype (Figure S3a). Chemotype 1 was dominated by limonene and sabinene, chemotype 2 by camphene, and chemotype 3 by β-terpinyl acetate and camphene. In stored root sesquiterpenoid profile, across all three chemotypes, β-sesquiphellandrene was the most dominant, followed by cis-β-farnesene (Figure S3b). However, the abundance of them varies between chemotypes.

### Stored root sesquiterpenoids are primarily affected by belowground herbivories

Belowground feeding by wireworms strongly increased the total concentration of stored root sesquiterpenoids (F = 75.25, *p* < 0.001; Table 1, Figure 1a) and stored root monoterpenoids (F = 14.64, p < 0.001; Table 1, Figure S4). Plant chemotype showed a significant main effect (F = 8.91, *p* < 0.001) on root sesquiterpenoids, and the chemotypic variation regulated the extent of the wireworm response with a strong two-way interaction (Chemotype × Worm: F = 11.56, p < 0.001), as the wireworms had stronger positive impacts in chemotypes 1 and 2 and less in chemotype 3 (Figure 1b). In contrast, aboveground feeding by aphids alone did not significantly affect stored root sesquiterpenoids, and the aphid and wireworm interaction were non-significant (Table 1). However, a significant three-way interaction of Chemotype × Aphid × Worm (F = 4.16, *p* = 0.022) and a marginal two-way interaction between aphid and chemotype (F = 3.01, *p* = 0.060) showed that the combined effect of wireworm & aphid on root sesquiterpenoids is likely chemotype-dependent. This pattern is indicated by the differences between wireworm & aphid impacts relative to control and aphid impacts across chemotypes, which was strongly increased in chemotype 1, but not affected in chemotype 2-3 (Figure 1b). We also tested the effect of naturally occurring biotic variables on the response variables. Powdery mildew was significant on stored root sesquiterpenoids (F = 5.84, p = 0.020; Figure S5), while thrips and *Coloradoa tanacetina* showed no significant effects (Table 1).

**Table 1:**
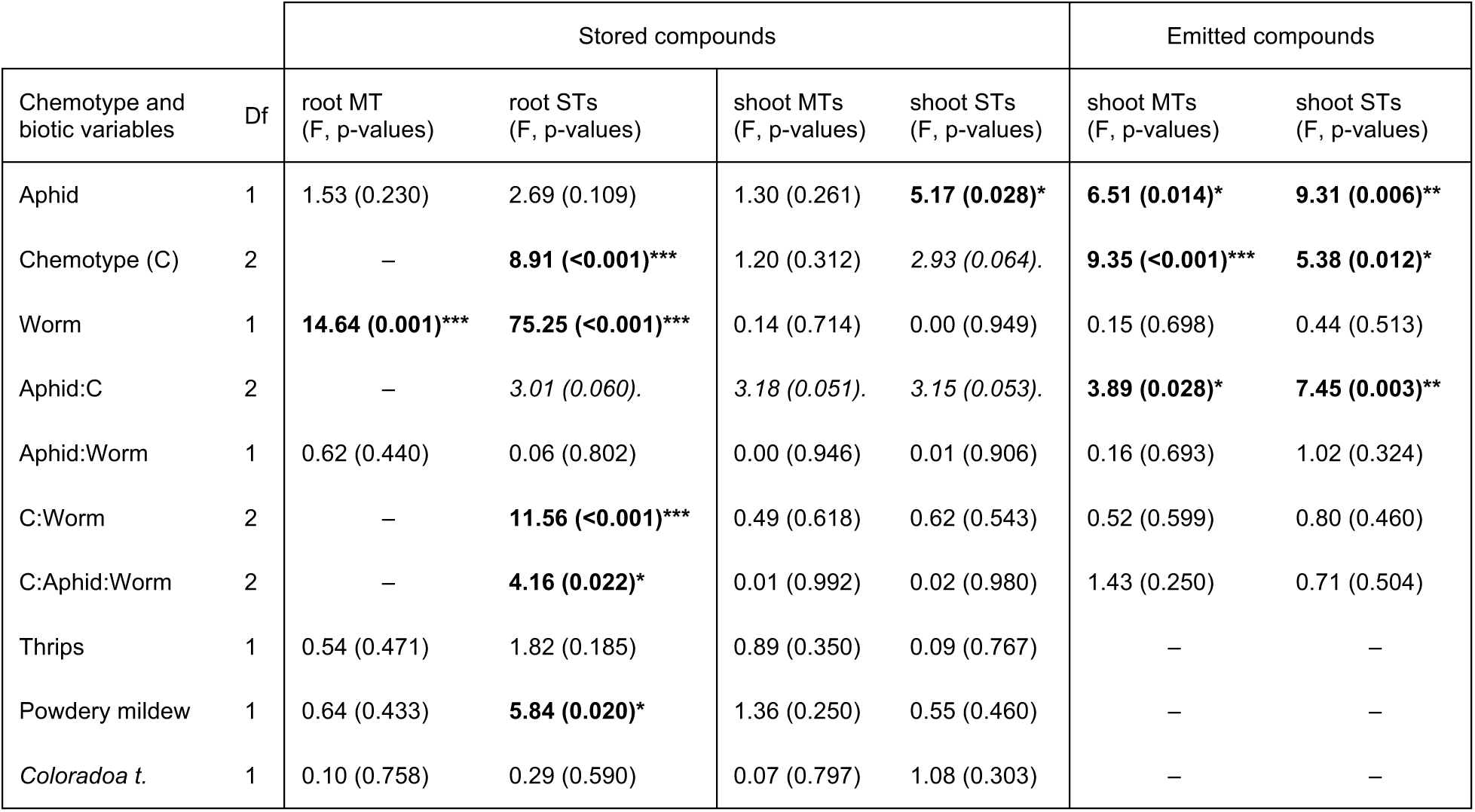
Analysis-of-variance summary for the linear model. The model depicts the effects of wireworm (Worm; *Agriotes spp.*), aphid *(Macrosiphoniella tanacetaria)*, chemotype, and unplanned biotic variables (thrips, powdery mildew, *Coloradoa tanacetina*) on stored (root and shoot) and emitted (shoot headspace) terpenoids. Response variables are stored root MTs and STs, stored shoot MTs and STs, and emitted shoot MTs and STs. Degrees of freedom (Df) and the F statistics with their associated p values are listed in the table. Significant values (p < 0.05) are highlighted in bold, and marginally significant values (0.05 < p < 0.1) in italics. P values smaller than 0.001 are reported as “<0.001”. In the stored-root MT analysis, we omitted the chemotype factor because excluding zero responses for the log transform left some chemotype * treatment combinations empty, making several coefficients aliased (non-estimable). For the emission models, thrips, powdery mildew, and *C. tanacetina* were excluded because headspace sampling occurred 10 days before the final harvest, whereas these covariates were assessed at harvest and thus including them would confound timing. Residual df by model: stored root MT = 21; stored root ST = 43; stored shoot MT = 44; stored shoot ST = 44; emitted shoot MT = 45; emitted shoot ST = 23.

Aphid had a significant main effect on the total concentration of stored sesquiterpenoids in leaves (F = 5.17, p = 0.028), but not on stored monoterpenoids. The Aphid × Chemotype interaction was borderline for both monoterpenoids (F = 3.18, p = 0.051) and sesquiterpenoids (F = 3.15, p = 0.053). Wireworm showed no main effects, and no interactions involving wireworm were significant for either terpenoid class in leaves.

### Chemotype-specific responses of root sesquiterpenoid *building blocks*

Pairwise correlations calculated for the 20 stored root sesquiterpenoids revealed strong associations between them (Figure 2b). Hierarchical clustering (Ward’s method) outlined four highly related groups, hereafter called building blocks, that are likely the products of farnesyl or nerolidyl cations in tansy (Figure 2a). Within each block, most pairwise coefficient values exceeded 0.7 (blue circles), supporting the proposal that each block represents distinct sesquiterpenoids (Figure S6) that are the products of a single enzymatic activity. Compounds highlighted by dashed frames in Figure 2a (e.g. modephene and petasitene) showed the strongest positive links to their proposed “parent” product, supporting the biosynthetic relationship. We used Sesquiterpenoid Synthase Database (bioinformatics.nl, Wageningen University & Research, see the full link in the references), an annotated survey of more than 300 plant sesquiterpenoids to show whether products originate from a farnesyl or nerolidyl cation. Block-wise responses of sesquiterpenoids to herbivores also strongly differed among chemotypes (Figure 3). In chemotype 1, wireworms consistently doubled the concentration of Block 1 (dominated by β-sesquiphellandrene) and Block 3 (dominated by α- and β-isocomene), and in Block 1, the dual attack enhanced the increase even further relative to control and aphid treatment. In chemotype 2, blocks 1, 2, and 3 showed a modest rise in wireworm groups, while in chemotype 3, none of the blocks responded to any herbivore treatment, mirroring its generally muted sesquiterpenoids plasticity.

**Figure 2:**
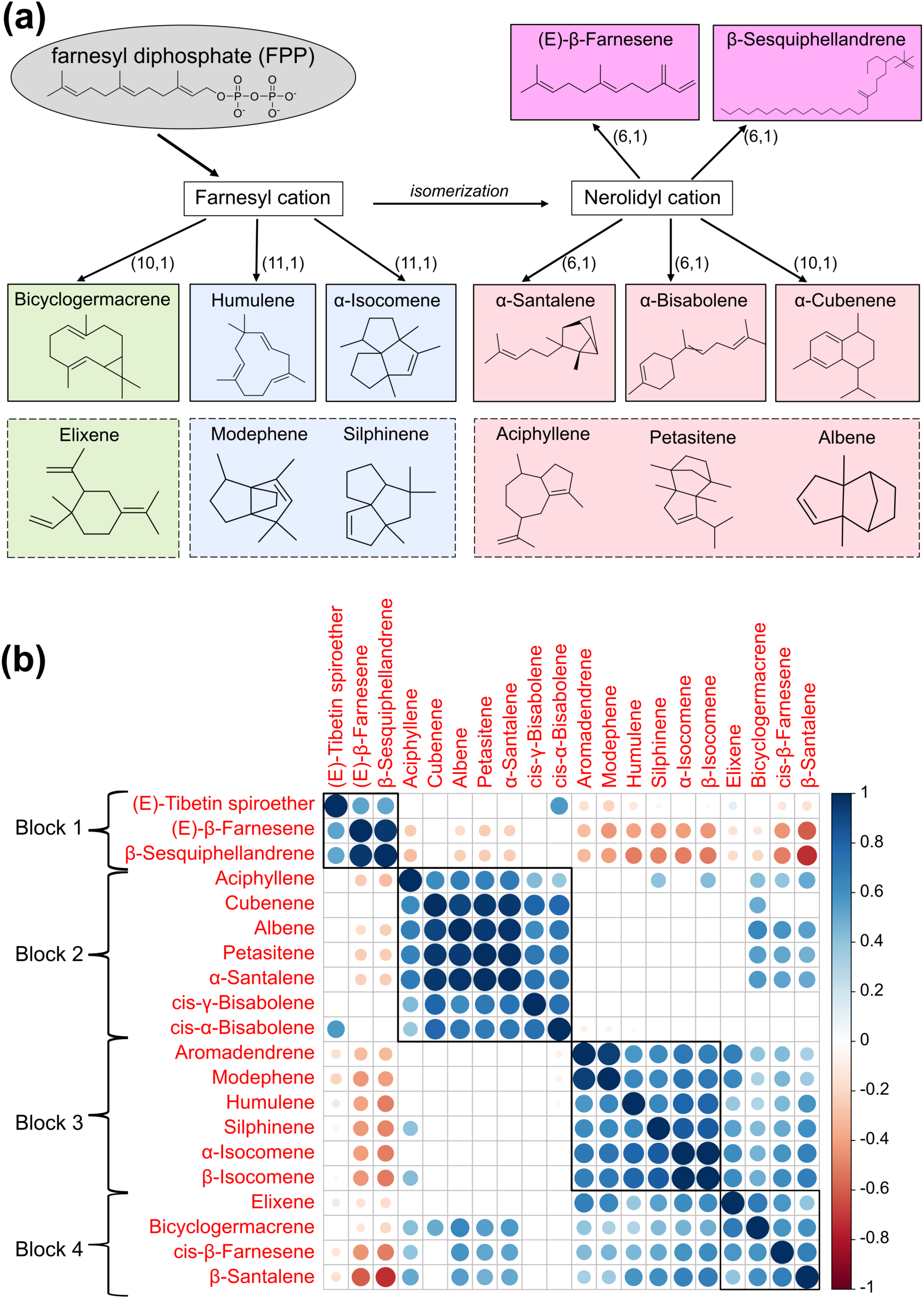
**(a)** Tansy root sesquiterpenoid classification scheme from the parent cations produced from farnesyl diphosphate. The numbers on top of each arrow show the cyclization from each cation. Four different colors show the corresponding sesquiterpenoids from different building blocks. The compounds highlighted with the dashed lines are proposed to show a biosynthetic relationship with the compound from the same pathway based on a strong correlation between them. The scheme was drawn using ChemDraw (version 23.1.2) **(b)** The correlogram showed pair-wise associations among the root sesquiterpenoids clustered in four building blocks, indicating a likely linked biosynthesis. Circle colors show the direction of the correlation: blue -positive, red - negative. Cells left blank correspond to non-significant correlations (p > 0.05). Black squares labeled Block 1 - Block 4 are the building blocks identified using hierarchical clustering.

**Figure 3:**
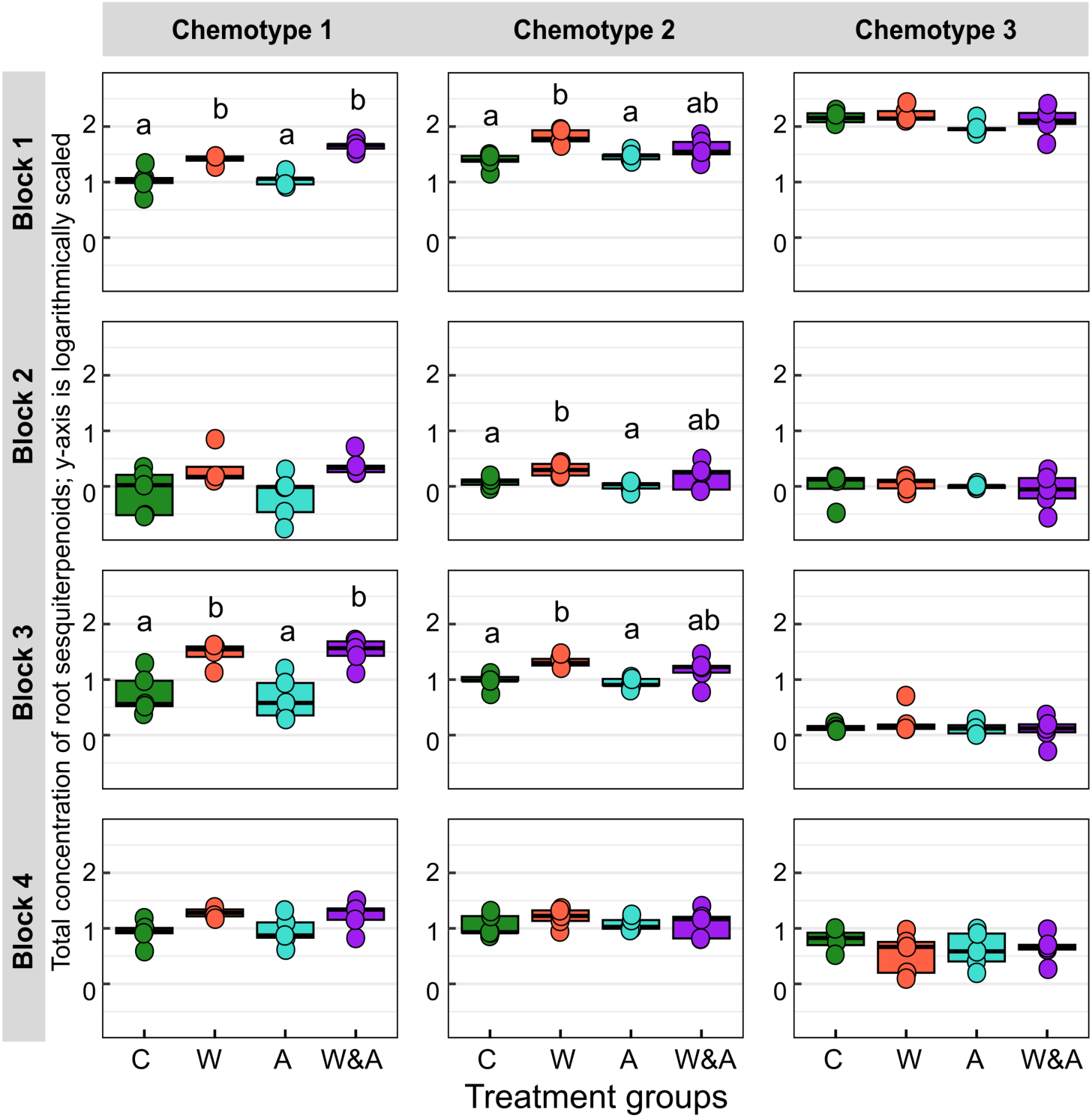
Block-wise response of stored root sesquiterpenoids to herbivore treatments across chemotypes. Blocks are defined based on the clustering of the root sesquiterpenoids shown in Figure 2b. Panels are arranged by block (rows 1–4) and chemotype (columns 1–3). Boxes show the interquartile range with medians; points are individual plants. Treatments on the x-axis: C = Control (green), W = Wireworm (red), A = Aphid (cyan), W&A = Wireworm & Aphid (purple). Within each panel, treatments were compared by one-way ANOVA (factor Treatment) followed by Tukey post-hoc tests. Letters show Tukey groupings: same letter = not significant (adjusted p ≥ 0.05); different letters = significant (adjusted p < 0.05). Panels without letters had a non-significant test and thus no pairwise groupings are shown.

### Volatile terpenoids of tansy showed minor shifts in leaves but not in roots

The monoterpenoids emission in leaves strongly increased in the presence of aphids (Aphid main effect: F = 6.51, p = 0.014, Table 1), also varying by chemotype (Chemotype main effect: F = 9.35, p < 0.001). The significant interaction between aphid and chemotype (F = 3.89, p = 0.028) showed that the emission of monoterpenoids in leaves was higher under aphid presence in chemotype 1, but not in chemotype 2-3 (Figure 4a). Similarly, the main effects of aphid, chemotype and their interaction term was significant on sesquiterpenoid emission (Figure 4b). However, these statistical terms should be interpreted cautiously, provided that sesquiterpenoid were detected in only 35 of 58 plants, yielding an unbalanced replication between control and aphid treated plants. Neither the wireworm main effect nor any interaction terms involving wireworm were significant.

**Figure 4.**
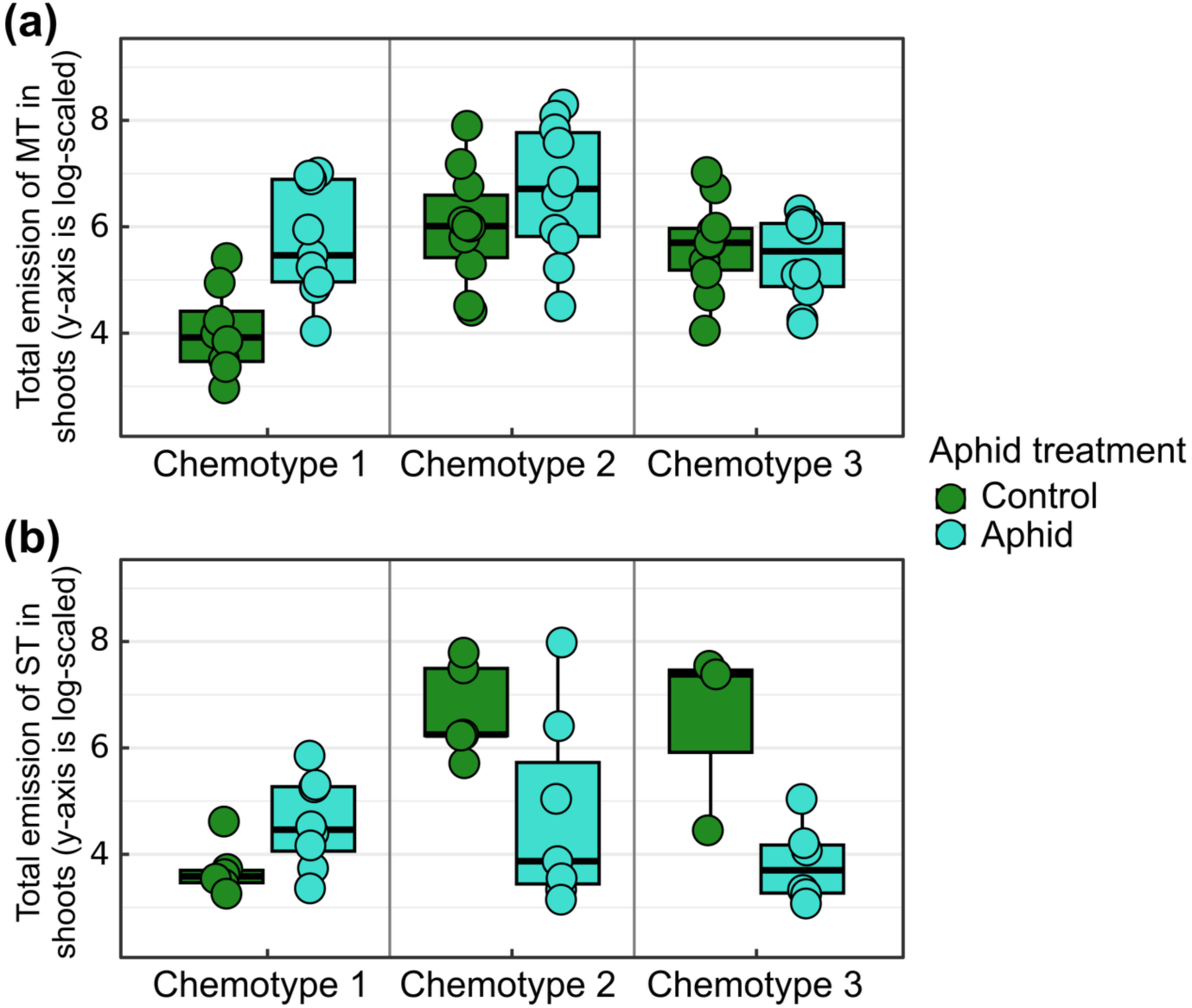
Boxplots of (a) total monoterpenoid (MT) and (b) total sesquiterpenoid (SQT) emissions (headspace analysis) from shoots under Control vs. Aphid treatments. Individual points show sample values and boxes span the interquartile range with horizontal median lines, and vertical grey lines separate chemotype blocks. Simple-effect contrasts of Aphid within each chemotype were obtained from the full model using *emmeans* averaging over Worm and p-values Bonferroni-adjusted across chemotypes. For monoterpenoids, chemotype 1 shows higher emissions under aphid (Aphid > Control; p = 0.002), chemotype 2, and 3 show no difference (p > 0.15). Sesquiterpenoid emission was statistically significant in chemotype 2 and 3 (p = 0.011; p = 0.005), but not in chemotype 1. The statistical tests for sesquiterpenoid emission should be interpreted with caution, as sesquiterpenoids were detected in only 35 of 58 plants. Note: the ‘Control’ legend pools plants where aphids are absent (Control + Worm; green), and the ‘Aphid’ legend pools plants where aphids are present (Aphid + Worm&Aphid; turquoise).

Root volatile emissions were generally low, but they were spatially structured within the rhizotron. The upper 3-5 cm (“top-zone”), where coarse roots are concentrated, emitted substantially more monoterpenoids than the lower-root zone (Bonferroni-adjusted Dunn tests: top- vs. low-zone p < 0.001; Figure. S7a). Despite a few outliers for sesquiterpenoids in the low-zone, the top-zone also showed higher sesquiterpenoid emission (p = 0.025; Figure S7b). Although the top-zone sampler clearly captured root-emitted mono- and sesquiterpenoids, the number of positive samples was limited (mono-: 25; sesquiterpenoids: 13), which precluded robust multivariable modeling of treatment and interaction effects. Accordingly, we present these root emissions descriptively and reserve formal hypothesis testing for a follow-up experiment with greater replication. Additionally, we visually examined tissue- and chemotype-specific patterns in emitted terpenoids on a heatmap. Samples clustered primarily by tissue, clearly separating leaves from roots (Figure. 5), consistent with expected differences between biosynthetic classes. Leaf emissions were dominated by monoterpenoids: 22 monoterpenoids were detected in leaves, whereas roots emitted only seven, four of which (α-terpinene, limonene, terpinolene, γ-terpinene) also occurred in leaves. By contrast, sesquiterpenoids were scarce in leaves but abundant and diverse in roots. Within each tissue, the emission bouquets varied with chemotype, broadly echoing the foliar chemotype classification seen in stored profiles.

**Figure 5:**
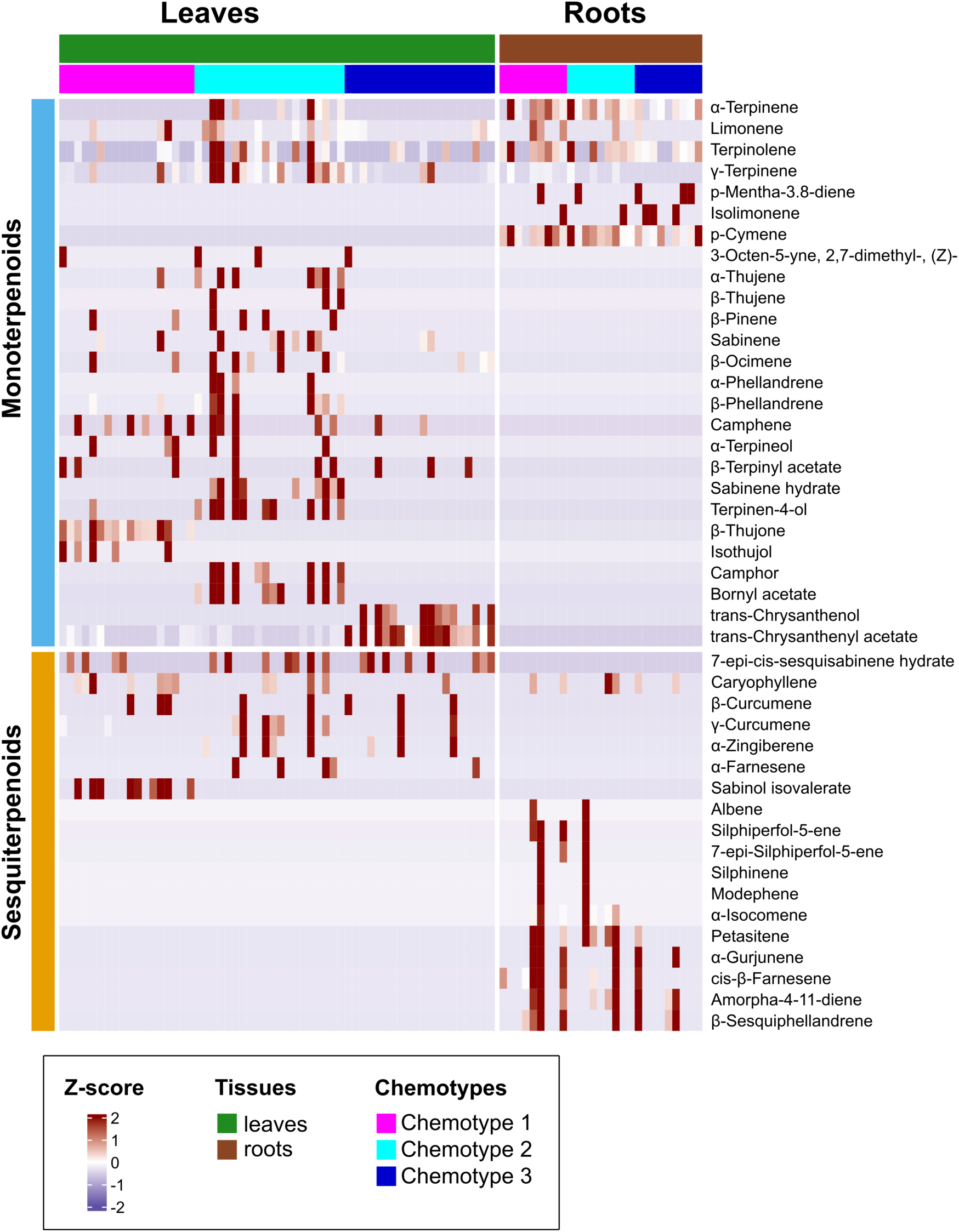
Heatmap of emitted terpenoid profiles in tansy leaf and root tissues across chemotypes, highlighting the patterns of tissue- and chemotype-specific and compound-class separation of tansy. Each column is an individual plant grouped first by leaf (in green bar) and root tissues (in brown bar) and by chemotypes: chemotype 1 in magenta, chemotype 2 in cyan, and chemotype 3 in blue. Monoterpenoids are displayed by sky blue bar in the top block, and sesquiterpenoids by gold bar in the bottom block. Cells show row-wise Z-scores: for each compound, values were log-transformed and standardized across all samples. Colors therefore represent relative abundance for that compound (red = above its mean; blue = below), not biological up/down-regulation. Because emissions are non-negative and some entries are zero, most below-average values cluster near zero (pale blue/white), whereas a subset of samples with higher emissions appears red.

### Chemotypes differ in root-system development and allocation dynamics

Time-course trajectories (Figure 6) demonstrated that root traits varied systematically among chemotypes throughout the monitoring period. Across all four metrics – root surface area, total root length, root volume and root-to-shoot ratio (in terms of root surface area / leaf area), chemotype 1 consistently produced the largest values, followed by chemotype 2 and 3 until around day 335 (12 days before the end of the experiment). Between days 320 and 330 (almost 9 weeks after the start of the experiment), chemotype 1 roots covered the highest values across traits (Figure 6). Over this interval, chemotype 2 reached about 80-90% and chemotype 3 about 70-80% of chemotype 1. All chemotypes followed a sigmoidal pattern but differed in growth. Chemotype 1 displayed the steepest rise until around day 330, and later reached its inflection point a few days earlier than chemotype 3 for surface area and length. Root volume, however, showed a substantial drop in chemotype 1 from around day 335, whereas volume in chemotype 3 maintained a continuous growth until the end of experiment. Chemotype 2 entered the saturation phase around the same time as chemotype 1. The reason chemotype 3 eventually exceeds the others in volume is because of its larger median root diameter (Figure S8), showing that this chemotype produced fewer but thicker roots. The root:shoot surface ratio peaked at about 1.7 in chemotype 1, 1.3 in chemotype 2, and 1.1 in chemotype 3 before all three converged toward ∼1.0 as leaf canopies expanded. Pairwise contrasts (Table S1) showed that chemotype 1 had higher values than the others for root surface area, total root length, and root:shoot ratio (χ², BH-adjusted p < 0.05). For root volume, chemotype 1 exceeded chemotype 2 (p = 0.027), while the contrast with chemotype 3 was not significant after adjustment (p = 0.055). Chemotypes 2 and 3 did not differ significantly for any trait (all p > 0.17). Furthermore, we observed a positive (R = 0.67, p = 0.01) correlation (Figure S9) between root surface area and the total concentration of stored root sesquiterpenoids within the pooled set of individual control plants that hinted at a synergetic relationship in the growth-defense system. The evidence is, however, insufficient as our sample size is too small to claim a definitive evolutionary synergy theory.

**Figure 6.**
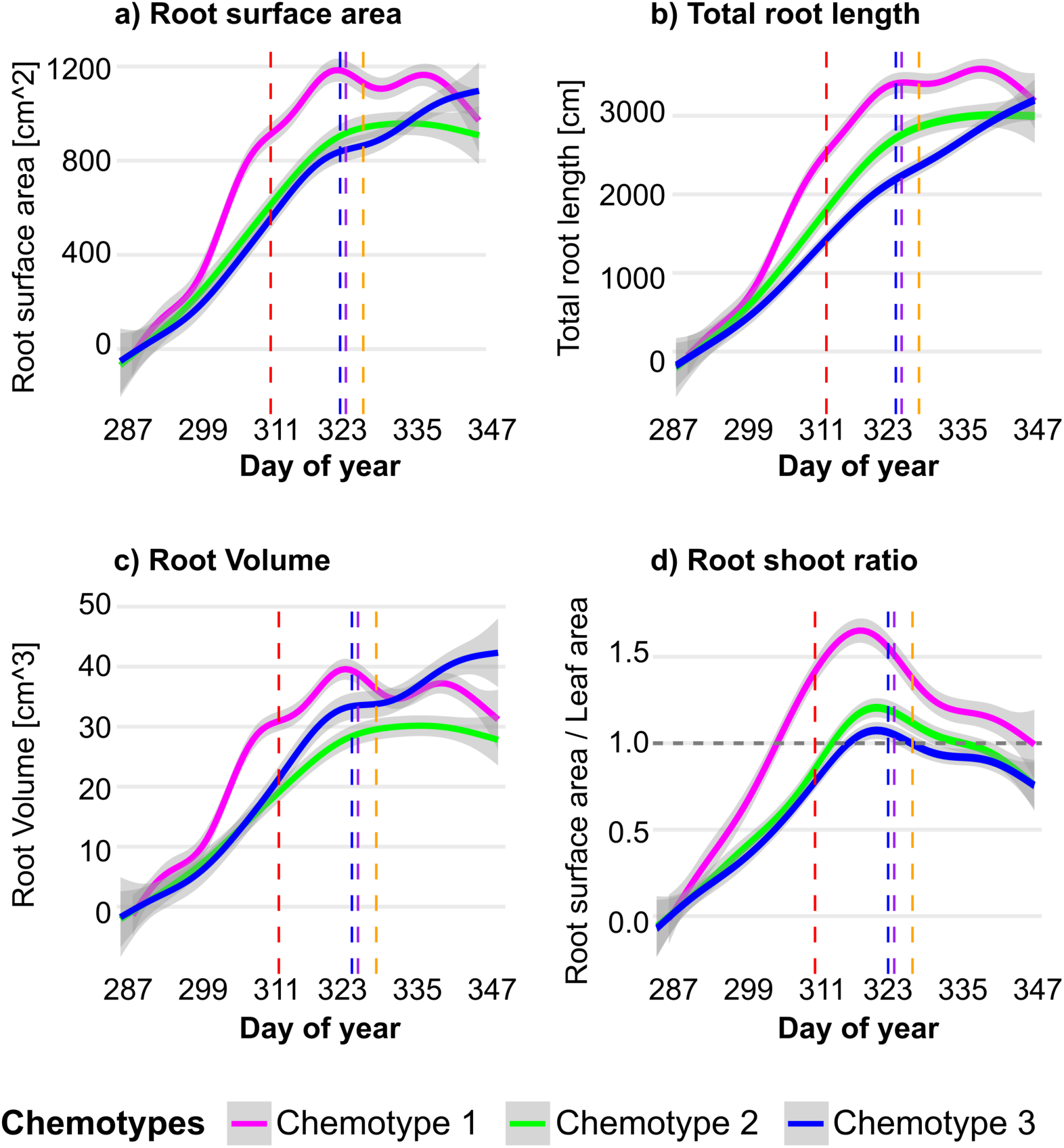
Growth dynamics of root traits by chemotypes. Curves are generalized additive model (GAM) smooths (thin-plate splines, *k* = 10; REML) fitted separately by chemotype, with 95% confidence ribbons. Panels show (a) root surface area, (b) total root length, (c) root volume, and (d) root:shoot ratio over the days of year (day 287–347). Magenta, green and blue lines correspond to chemotype 1, 2, and 3, respectively. The vertical dashed lines depict different events that occurred during the experiment; from left to right: the start of wireworm treatment (day 313), and of aphid treatment (day 325), twister analysis in the rhizotrons (day 326), initial harvesting (day 329), respectively. Note: the initial harvesting was conducted for transcriptomic analysis, which is not part of the current study. Chemotype 1 exhibits the most rapid early root expansion and reaches its maximum root:shoot ratio around day 320.

## Discussion

In this study, we demonstrate that above- and belowground herbivory in *Tanacetum vulgare* elicits compartment-specific responses that reveal localized and specific defense strategies. Feeding by a belowground root herbivore strongly induced production of mono- and sesquiterpenoids in the roots, whereas feeding by an aboveground phloem-feeding herbivore induced predominantly volatile monoterpenoid emissions. Using different chemotype lineages, we reveal that there are both generalizable and chemotype-specific responses to herbivory. We identify strong biosynthetic clustering highlighting terpenoid biosynthesis pathways, which are distinctly impacted by herbivory. Our findings provide strong evidence for the specific ecological roles of mono- and sesquiterpenoids and associated biosynthetic machinery.

Belowground wireworm feeding elevated the total concentration of stored root terpenoids, while having no influence on shoot pools or emissions. Additionally our results showed that aphids alone did not affect root terpenoids. Yet in combination with wireworms, aphids amplified the sesquiterpenoid increase in chemotype 1, and had no detectable effect in chemotype 2 and 3. Thus, our study suggests that phloem feeders do not elicit systemic root induction on their own, but can modify root defenses triggered by belowground herbivores in a chemotype-dependent manner. A comparable pattern was reported by Bezemer et al. (2004) who demonstrated that terpenoid-aldehydes increased markedly in roots of *Gossypium herbaceum* seedlings after wireworm *(Agriotes lineatus)* feeding. While the authors demonstrated a slight systemic signal reaching the leaves, the quantitatively stronger reaction was contained in the roots. This is not at odds with the broader above- and belowground literature. Generally systemic outcomes depend strongly on the inducing guild and hormone cross-talk. For instance, phloem feeders typically trigger salicylic acid (SA) and can antagonize jasmonic acid (JA) signaling (Erb & Reymond, 2019), whereas many sesquiterpenoid pathways, including root sesquiterpenoids are JA-dependent (Tholl, 2015). Within this framework, the lack of general aphid-induced responses in root sesquiterpenoids aligns with SA-JA cross-talk. In a related study, Karssemeijer et al. (2020) showed that phloem-feeding cabbage aphids *(Brevicoryne brassicae)* did not activate the root JA pathway, whereas chewing caterpillars *(Plutella xylostella)* did, and this translated into reduced performance of the root herbivore *Delia radicum*. The authors interpret this as guild-specific systemic cross-talk (SA-biased aphids vs JA-biased chewers), consistent with our lack of JA-dependent root sesquiterpenoid induction under aphid feeding. By contrast, systemic shoot-to-root responses can occur in other systems. For instance, in Arabidopsis, aboveground aphid feeding *(Brevicoryne brassicae)* triggered systemic transcriptomic changes in roots, whereas root attack by *Heterodera schachtii* did not produce systemic changes in shoots (Kutyniok et al., 2014). The authors interpret these patterns as attacker-specific, signaling-mediated responses under low infestation, consistent with SA-JA cross-talk. In their earlier metabolomics work they described ‘minute’ systemic metabolic shifts under similar conditions (Kutyniok & Müller, 2012). Taken together, our data indicate that while aphids do not directly induce JA-dependent root sesquiterpenoids, they can modulate root defenses during cross-guild attack in a chemotype-specific manner.

Furthermore, our findings demonstrated a pronounced chemotype-specific induction of root sesquiterpenoids following a wireworm attack. Chemotypes 1 and 2 induced their sesquiterpenoid pools under wireworms, and the combined effect of wireworm & aphid further boosted the induction in the root chemotype 1. In contrast, chemotype 3 remained mostly unchanged. This highly chemotype-specific response is very likely linked to building blocks of sesquiterpenoids that we identified using a hierarchical clustering method. These blocks are proposed to reflect a biochemical signature commonly observed when a single terpenoid synthase (TPS) produces a diverse array of products from the farnesyl diphosphate substrate. Upon evaluating the response of each proposed block to each treatment within each chemotype, we found a significant induction effect of Block 1 (defined by β-sesquiphellandrene) and Block 3 (defined by α- and β-isocomene) triggered by wireworms in chemotype 1, thereby supporting the remarkable plasticity of this chemotype that we observed before. This block-level and chemotype-specific induction of root sesquiterpenoids might be linked to discrete modules of sesquiterpenoids synthesized from a single enzyme. For instance, in spotted knapweed *(Centaurea stoebe),* two root-specific synthases, CsTPS4 and CsTPS5, reproduce the whole sesquiterpenoid profile, indicating that a single enzyme could regulate an entire chemical module (Gfeller et al., 2019). Similarly, in maize, the inducible β-caryophyllene synthase TPS23 is expressed in ancestral and European lines but is transcriptionally absent in the majority of North American types, along with the entire inducible volatile module (Köllner et al., 2008). By analogy, the strong induction of Blocks 1 and 3 in chemotype 1 suggests that this chemotype possesses active alleles of the TPS genes for those blocks, whereas the other chemotypes probably carry silenced or low-expression variants. Leaf transcriptomics in tansy (Clancy et al., 2020) showed broad TPS expression, but few herbivory-responsive genes: a putative (–)-germacrene D synthase (TanvuEGr017925) and two (E)-β-farnesene-like TPSs (TanvuEGr029614, TanvuEGr007220). TanvuEGr029614 was strongly induced by aphids, while TanvuEGr007220 increased under combined aphid & caterpillar attack. These leaf data support inducible, chemotype-specific TPS control of volatile emissions. Because β-sesquiphellandrene and isocomene dominate tansy roots, analogous root TPSs likely exist, but remain uncharacterized. Because we used statistical methods alone to define these blocks, a crucial next step would require validating them via transcriptomic profiling and heterologous expression of candidate TPS genes in tansy roots.

Following 14 days of aphid feeding and 36 hours of volatile collection using twisters, headspace analysis of leaf samples indicated a significant increase in shoot monoterpenoid emissions in chemotype 1 (β-thujone dominated in leaves) while chemotype 2 (camphor and sabinene-hydrate dominated), and 3 (trans-chrysanthenyl-acetate dominated) exhibited no variation, suggesting that the emission of monoterpenoids under aphid attack is specific to chemotypes. Solvent extracts of the same leaves demonstrated only weak alteration in stored monoterpenoid pools, a pattern that aligns with the kinetic model in which sap-feeding insects primarily stimulate terpenoid-synthase genes, resulting in a rapid *de novo* “emission pool” while the slow turnover reservoirs in glandular trichomes remain mostly unaffected (Clancy et al., 2020; Niinemets et al., 2013). Why is the emission of monoterpenoids favored? Monoterpenoids generally have higher vapor pressures than sesquiterpenoids, allowing them to diffuse out easily through the cuticle, whereas sesquiterpenoids, having higher boiling points and lower vapor pressures, are often retained in storage pools and are emitted slowly from intact leaves (Mofikoya et al., 2019). These physicochemical differences help explain why aphid feeding significantly enhanced monoterpenoid emissions compared to sesquiterpenoids, given the importance of enzymatic regulation in *de novo* synthesis (Niinemets et al., 2013). Clancy et al. (2020) likewise documented analogous storage versus emission decoupling and further revealed that aphid pre-infestation amplified the caterpillar-induced monoterpenoid spike in certain tansy chemotypes, but not in others. Our findings indicate that phloem-feeding aphids induce a chemotype-specific, *de novo* release of readily diffusive terpenoids, while primarily preserving the storage pools. We additionally collected the terpenoids from roots and found that the tissue-specific partitioning of emitted terpenoids parallels our earlier findings on stored pools (Rahimova et al., 2025), where mono and sesquiterpenoid profiles were clearly localized between shoots and roots. Despite a substantial number of replications, mono- and sesquiterpenoids were detected in only a few root samples, limiting our ability to statistically assess herbivory impacts. To effectively evaluate whether wireworms alter root terpenoid emissions, a specific, root-emission-focused methodology is required. For instance, Lee Díaz et al. (2022) used a sterile two-compartment Petri-dish and passively sampled root VOCs in situ from tomato seedlings 24 h after *Spodoptera exigua* attack, using HiSorb/PDMS sorbents and TD-GC-QTOF-MS. In addition to demonstrating an in situ capture setup, they indicated that root volatiles vary under herbivory and that the choice of methodology (HiSorb vs PDMS) influences the observed outcomes, advocating for multi-sorbent sampling.

Our results showed that chemotype identity is linked to not only terpenoid chemistry but also to root architecture. Chemotype 1 maintained the largest root surface area and length and the highest root:shoot ratio across the pre-harvest window. Chemotype 2 followed a similar rise but leveled off earlier at lower values. Chemotype 3 had the smallest surface area and length yet formed comparatively thicker roots (higher median diameter) and thus produced the larger root volume towards the end of the experiment. Because in tansy, coarse roots contain a higher total sesquiterpenoid concentration than fine roots (Rahimova et al. 2025), a greater fraction of coarse roots in chemotype 3 provides the plausible explanation for its higher constitutive root sesquiterpenoid storage that we observed in the present study (Figure 2b, control boxes). Additionally, a positive linkage between total root sesquiterpenoids and surface area suggests a beneficial association between root growth capacity and inducible defense, rather than a classic growth-defense trade-off. A comparable *“defense-supports-growth”* effect has been documented in the *Populus-Laccaria* symbiosis, where young poplar trees associate with the ectomycorrhizal fungus *Laccaria bicolor.* The fungus releases a bouquet of volatile sesquiterpenoids dominated by (-)-thujopsene that acts as an airborne signal to the plant. Exposure to this sesquiterpenoid blend reprogrammed the poplar’s root system, increasing the number and length of lateral roots and greatly expanding root surface area (Ditengou et al., 2015). Together with prior studies, our results suggest that chemotype co-varies with morphological traits, and observed ‘chemotype effects’ could stem from terpenoid chemistry itself or from other co-varying traits. Distinguishing these alternatives will require targeted causal tests to investigate this putative growth and sesquiterpenoid induced defense theory.

## Conclusions

In conclusion, the present study demonstrated that tansy exhibits highly compartmentalized, chemotype-dependent defense strategies. Wireworm feeding triggers a robust response in the mono and sesquiterpenoid concentrations in the roots yet fails to deliver any measurable upward signal to the shoots. Whereas aphid attack on leaves elicits a de-novo burst of monoterpenoid emissions without altering root chemistry. Among the three chemotypes, chemotype 1 launched the strongest inducible defense both in shoot and root tissues and produced larger root systems and higher root:shoot ratio. Collectively, these patterns indicate that tansy allocates its inducible chemistry and root architecture among tissues in a manner that depends on both the attacker and the plant’s identity. The subsequent crucial phase is to conduct a high-quality *Tanacetum vulgare* transcriptomic analysis. Such a platform will allow us to identify terpenoid synthase (TPS) genes that underlie each sesquiterpenoid “block”, and link genetic variation in TPS modules to both inducible chemistry and root system architecture.

## Acknowledgments

The authors are truly grateful to the EUS team, particularly to Ulrich Junghans, Petra Seibel, and Mustafa Özden in Helmholtz Munich, and Sarah Sturm in TUM, for their continuous support throughout the experiment and final harvesting.

## Contribution of authors

J-P.Schnitzler, R.Heinen and H.Newrzella designed the experiment; H.Newrzella gathered and evaluated the raw data, performed the statistical models, drafted the figures and the manuscript with the guidance from JP.Schnitzler, R.Heinen and W.Weisser; G.Gerl, P.Kary, and A.Sigalas set-up the experiment in the greenhouse; R.Heinen, A.Neuhaus, and L.Ojeda-Prieto propagated the plants; H.Newrzella, CM.Vilasenta, G.Gerl, P.Kary, R.Heinen, A.Neuhaus, and L.Ojeda-Prieto; CM.Vilasenta further analyzed and extracted the root images and features with the supervision of JB.Winkler; H.Newrzella and I.Zimmer conducted the leaf and root extraction analysis; B.Weber performed GC-MS runs and calibration analysis; J-P.Schnitzler and JB.Winkler supervised the 60 day of experiment in the greenhouse.

## Data availability

All data (GC–MS tables for leaf and root tissues, volatile emission matrices, imaging-derived root traits (daily time series), and statistical R scripts) will be deposited publicly in Dryad with a DOI upon acceptance.

## Supplementary Figure and Table Legends

**Figure S1:**
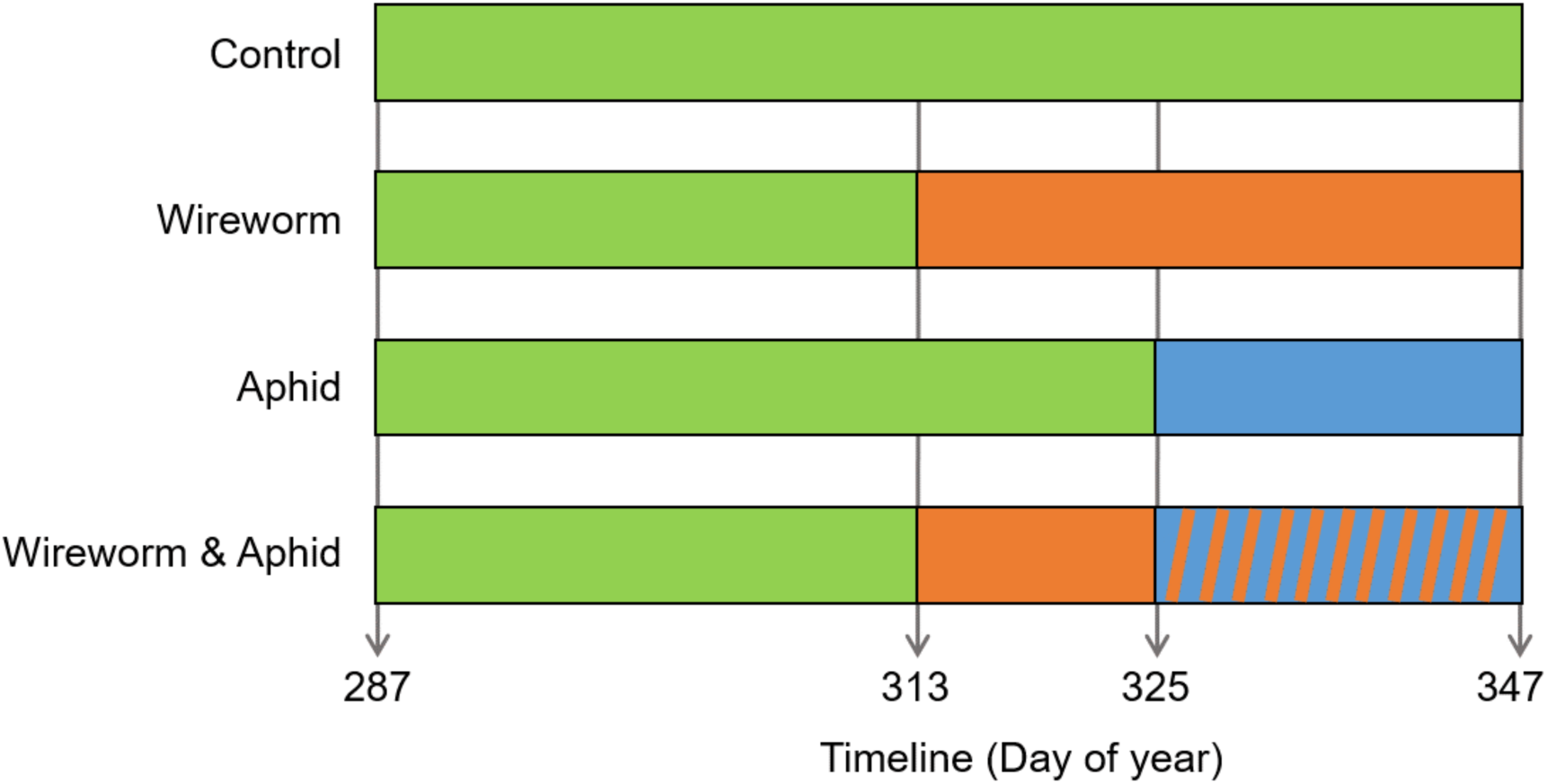
Treatment timeline of the tansy greenhouse experiment. Each horizontal bar represents one treatment: an unmanipulated control, wireworm only, aphid only, and wireworm & aphid groups. During the initial stabilization phase of the plants, all plants were maintained under identical conditions. Green segments show the establishment period (Day of year, 285–313), orange segments the wireworm phase (started on day 313), and blue segments the aphid phase (started on day 325). The experiment ended on day 347.

**Figure S2:**
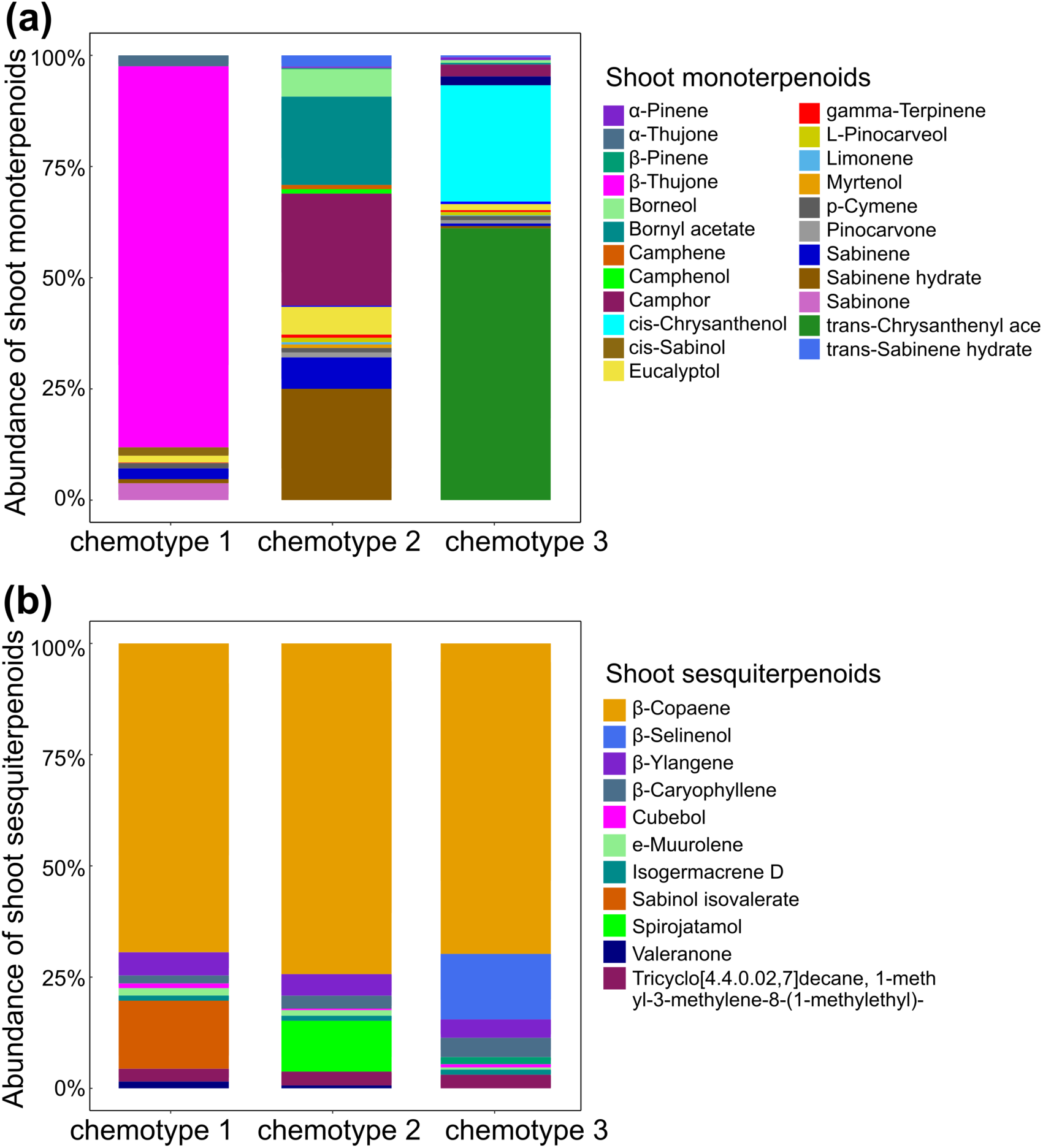
Terpenoid fingerprints of three chemotypes of tansy. Stored monoterpenoids (a) and sesquiterpenoids (b) in shoots. Each stacked column represents a distinct chemotype (1-3), with segment height illustrating the relative abundance of the compounds depicted in the accompanying legends.

**Figure S3.**
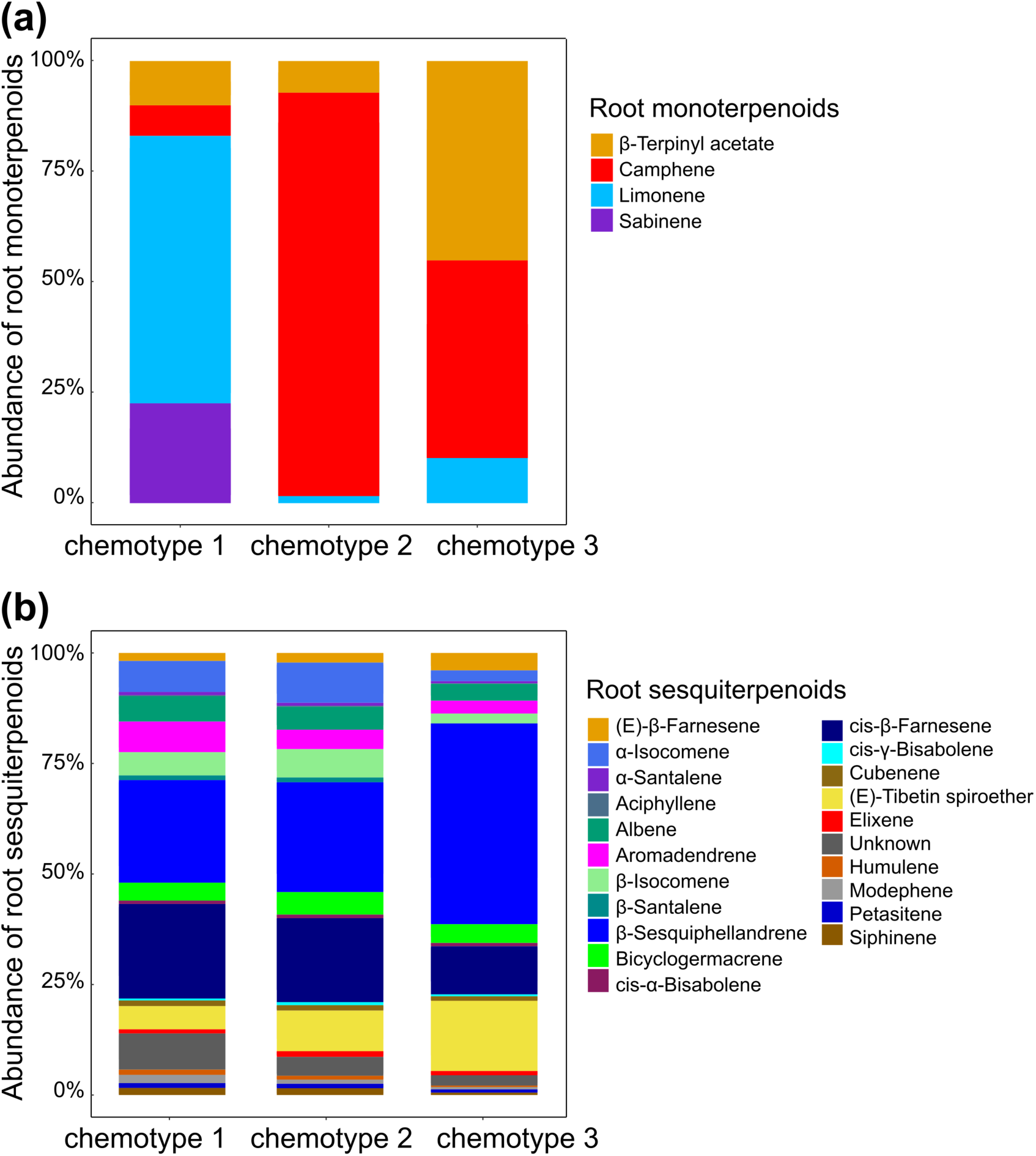
Root terpenoid profiles of three tansy chemotypes. (a) Stored monoterpenoids; (b) stored sesquiterpenoids. Each stacked bar represents chemotypes 1-3, and segment heights indicate the relative abundance of the compounds listed in the legends.

**Figure S4:**
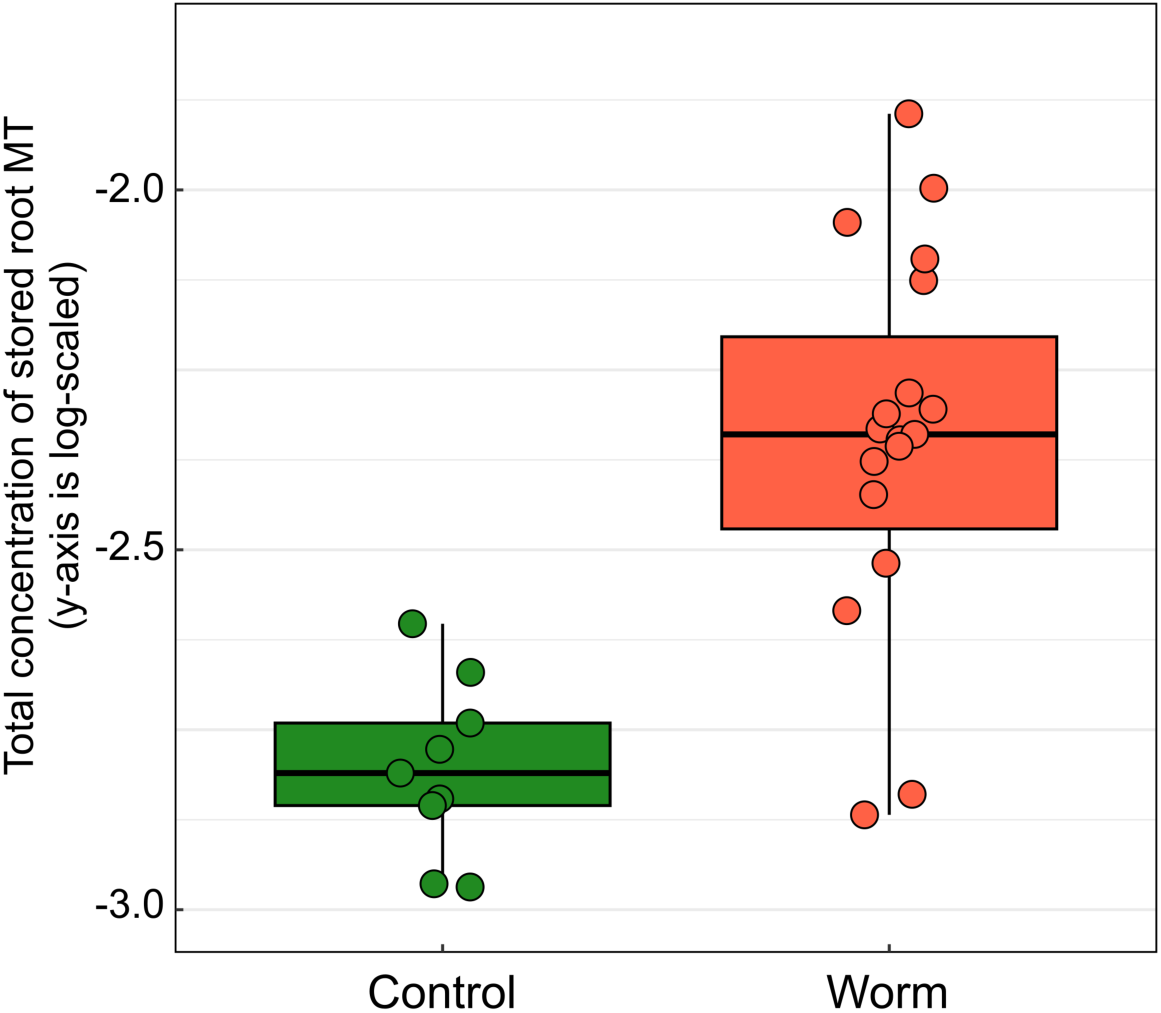
Boxplots of the log-transformed total concentration of stored root monoterpenoids (MT) in Control vs Worm-treated plants. Worm-treated roots had higher monoterpenoids than controls (Type III ANOVA on full model: Worm effect F = 14.62, p = 0.001).

**Figure S5:**
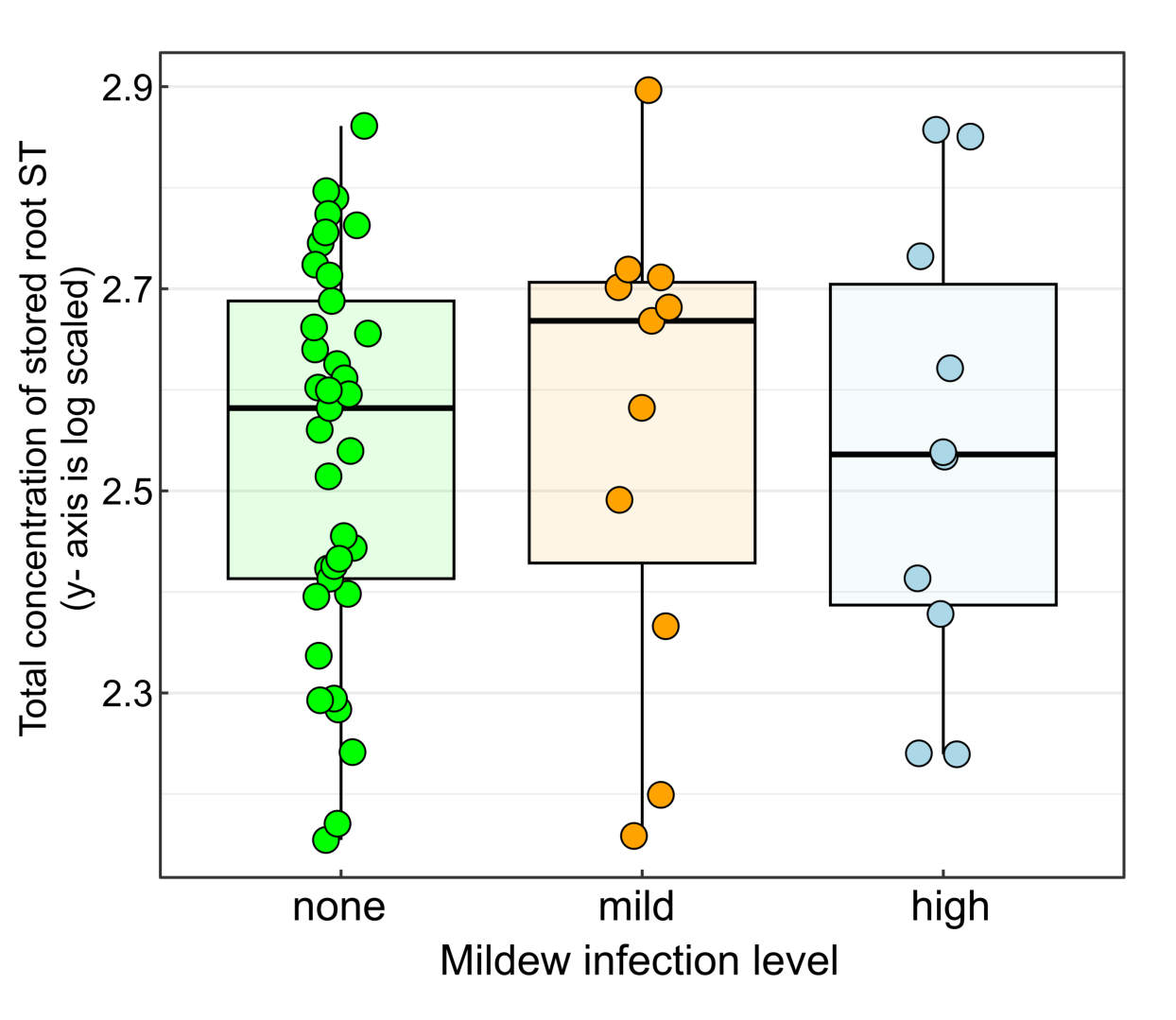
Distribution of total concentration of stored root sesquiterpenoids (ST) across mildew infection levels. Boxplots show values (y-axis on a log scale) for plants with none, mild and high mildew infection.

**Figure S6.**
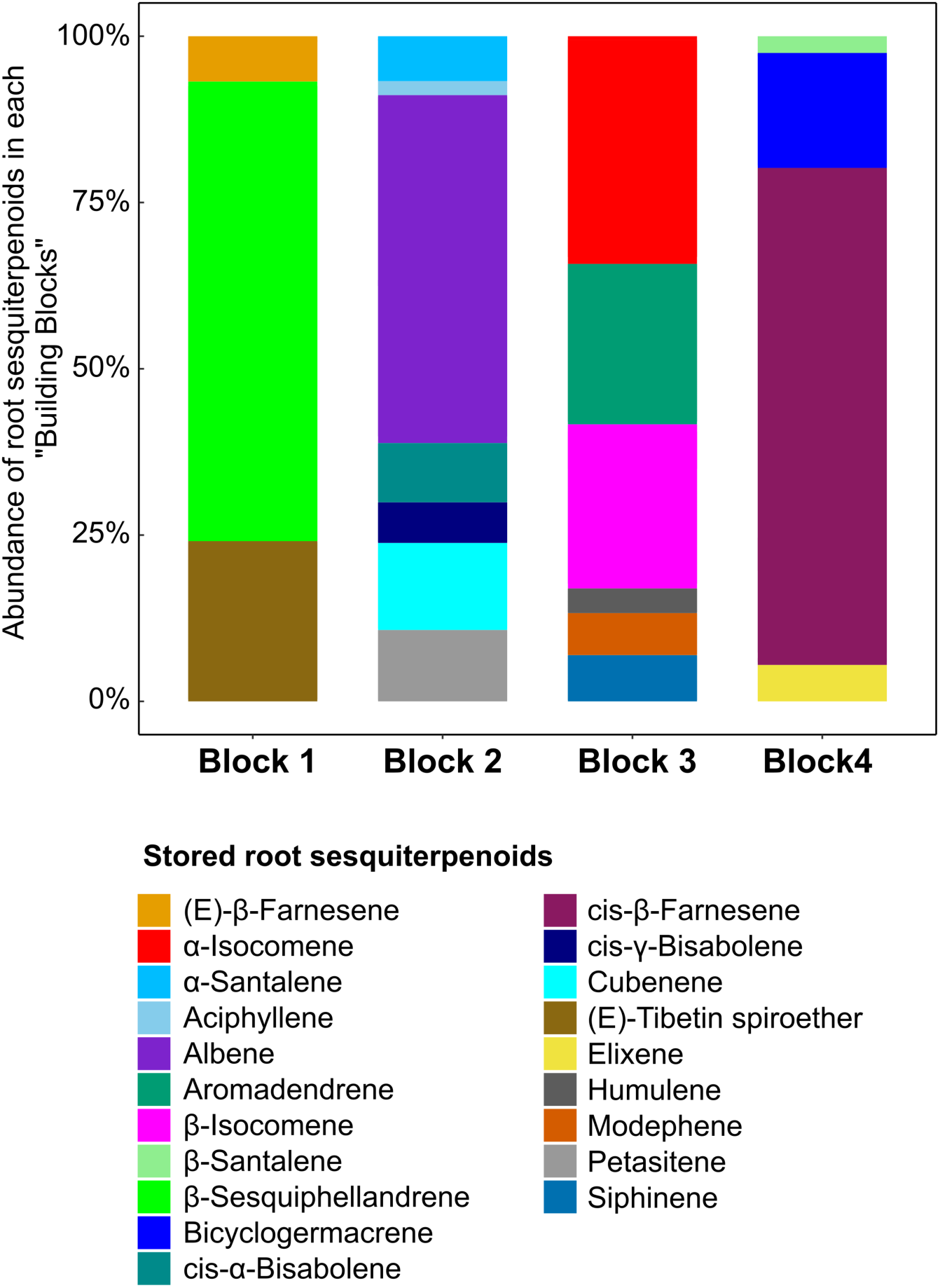
Relative abundance of stored root sesquiterpenoids in each building block. Block 1 is mostly dominated by β-sesquiphellandrene, Block 2 by albene, Block 3 by α- and β-isocomene, and Block 4 by cis-β-farnesene.

**Figure S7.**
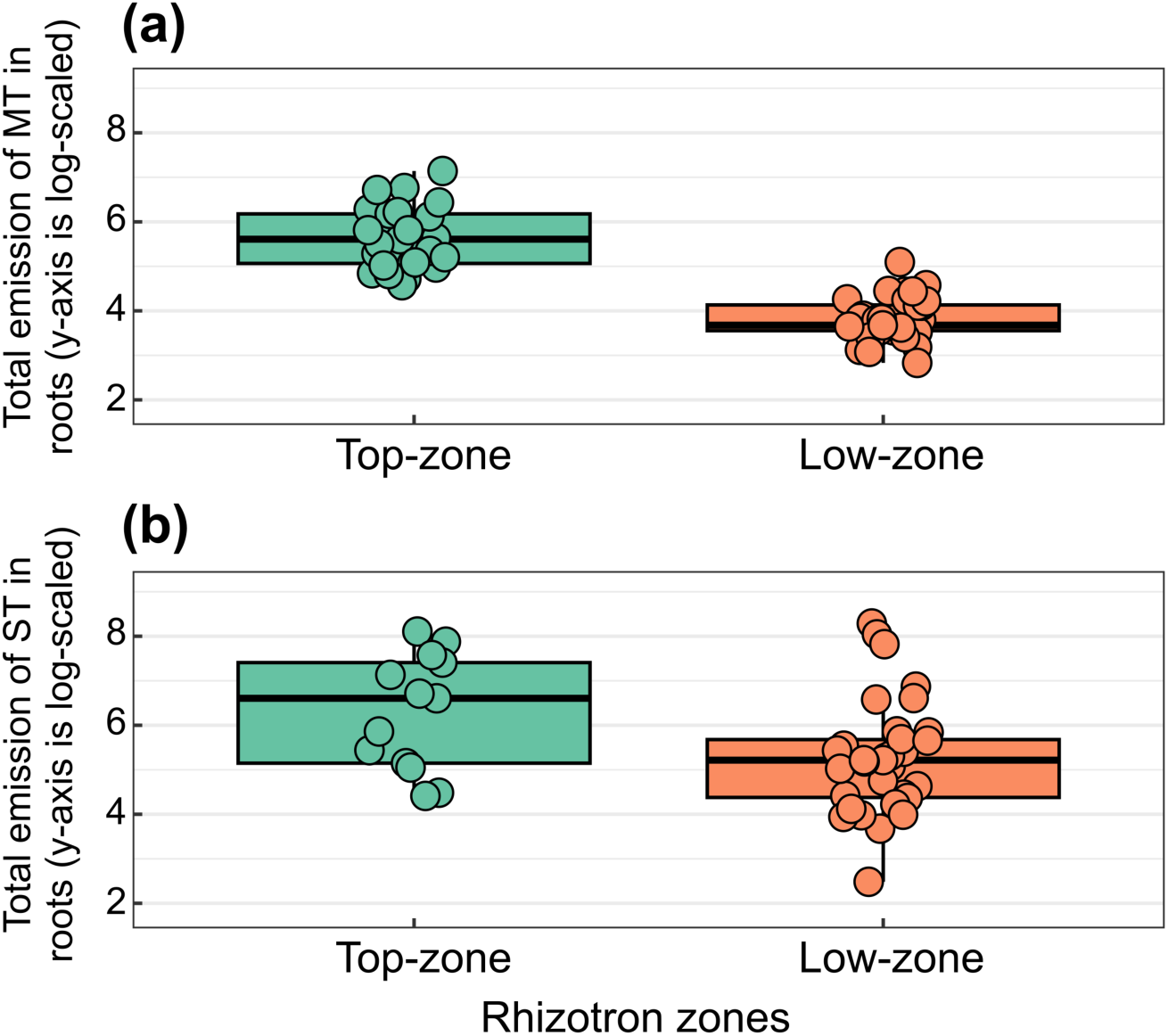
Root (a) monoterpenoid, and (b) sesquiterpenoid emissions across rhizotron zones: 1) top-zone – twister ∼5 cm below the surface (near coarse roots); 2) low-zone – twisters on individual lower roots (30-40 cm from the top) enclosed with foil bags. The soil-background check showed no terpenoid peaks and was t herefore omitted from the figure and analysis. Bonferroni-adjusted Dunn tests showed higher monoterpenoid and sesquiterpenoid emission in the top-zone relative to the low-zone (p < 0.001; p = 0.025).

**Figure S8.**
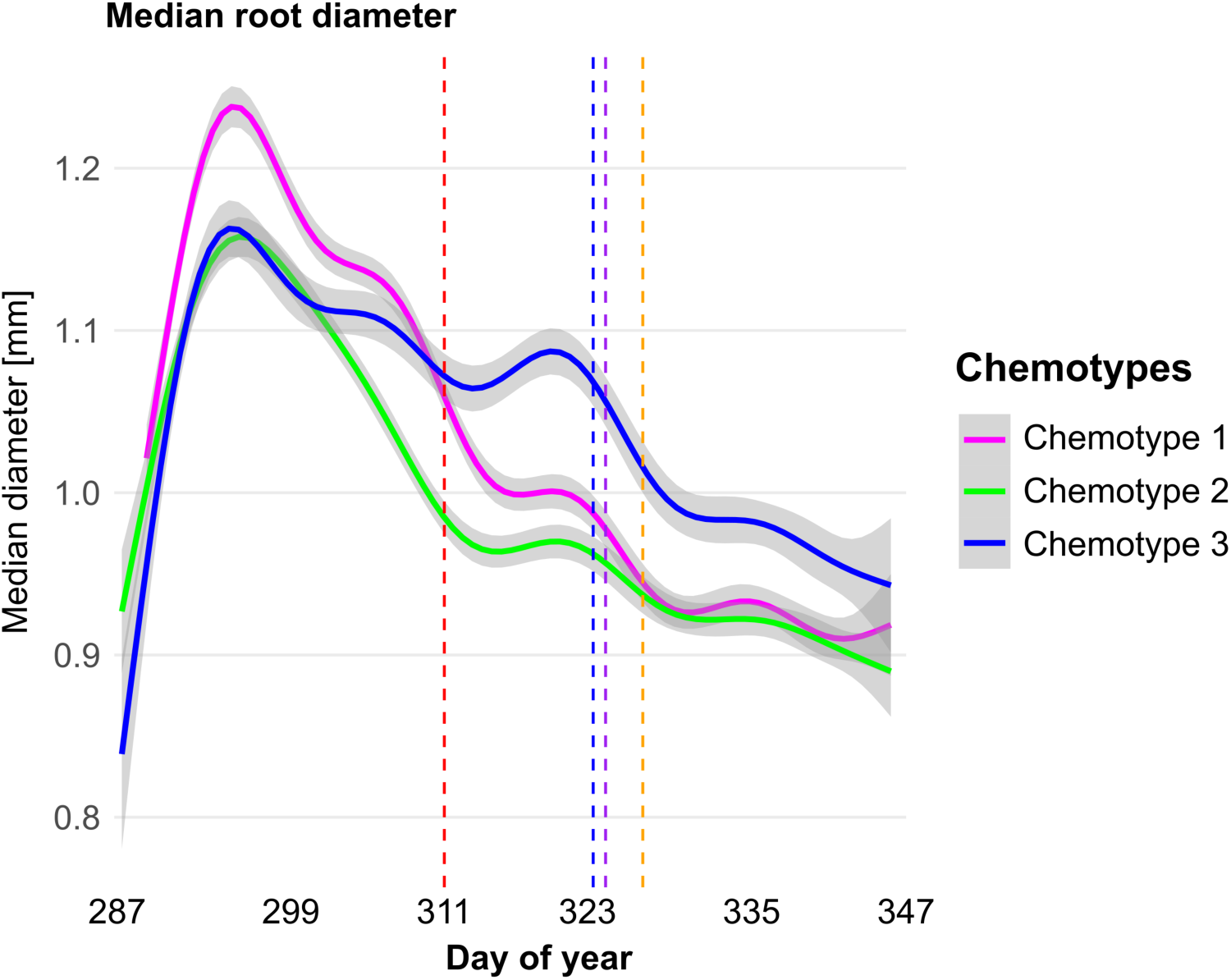
Median root diameter over the days of year (DOY 287–347) differed by chemotypes. Magenta, green, and blue curves show chemotype 1, chemotype 2, and chemotype 3. Vertical markers show, from left to right, the onset of wireworm treatment (day 313), aphid treatment (day 325), the rhizotron twister analysis (day 326), and the initial harvest (day 329).

**Figure S9.**
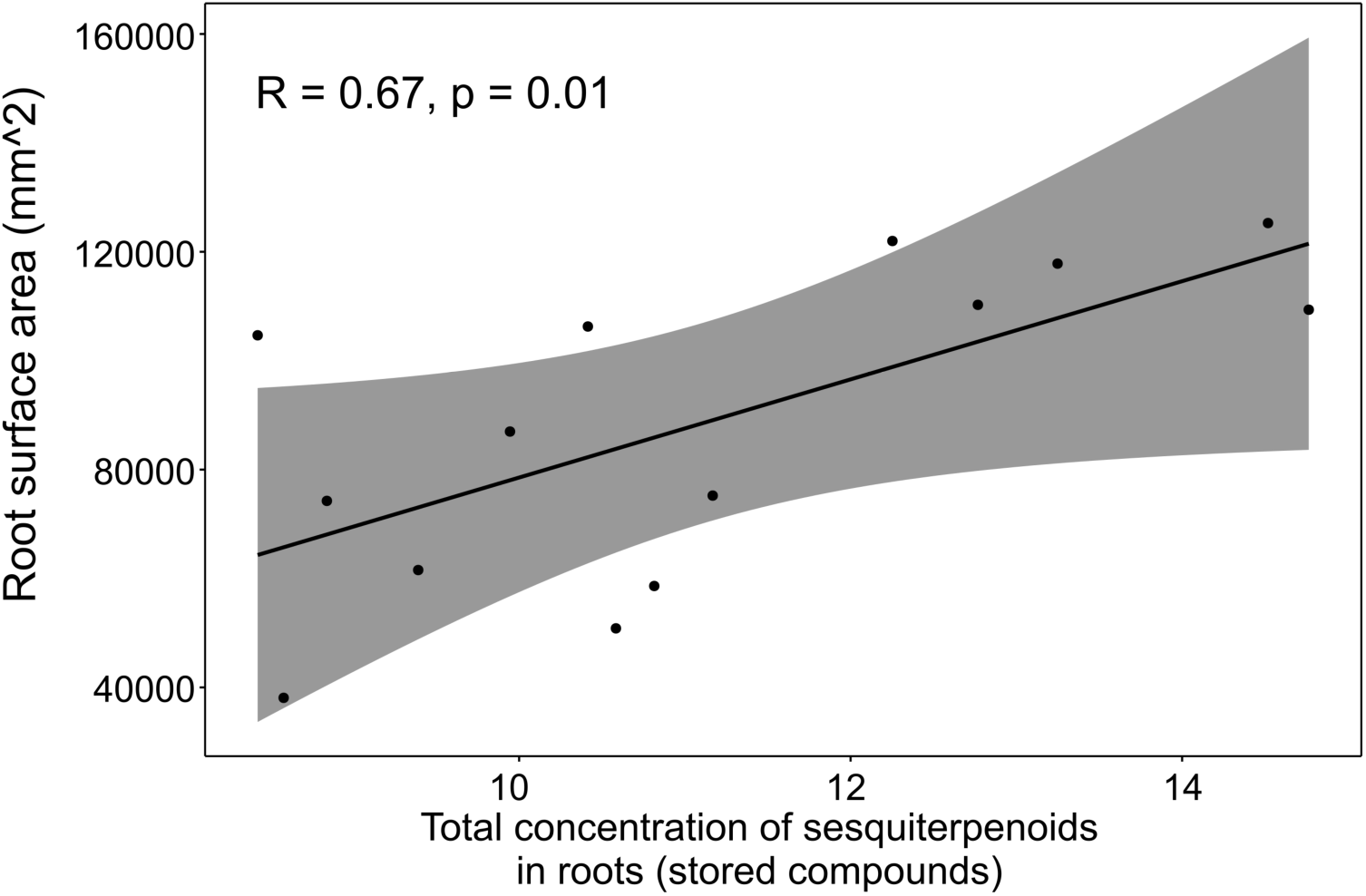
Root surface area (mm²) measured a week before the final harvest is plotted against the total concentration of stored sesquiterpenoids (Pearson’s R = 0.67, p = 0.01) in the roots of untreated plants (n = 14). The solid line shows the fitted linear regression with the 95% confidence interval (grey band).

**Table S1.** Pairwise chemotype contrasts for root traits from generalized additive mixed models (GAMMs). Models were fit with mgcv using REML and included chemotype-specific smooths of day-of-year (factor-smooth, *k* = 10) and a plant-level random intercept. Estimated marginal means were averaged uniformly over the observed time range. Pairwise differences were tested with Wald χ² (df = 1) from the EMM contrasts, and p-values were Benjamini-Hochberg-adjusted within trait. We limited statistical inference to data collected between day 287 and 329 (from the first measurement until the initial harvest) as this time window provided the most complete, uninterrupted observations.

| **Response** | **Comparison** | **Wald $\chi^2$** | **BH-adjusted p** |
| --- | --- | --- | --- |
| Surface area | Chemotype 1 – Chemotype 2 | **5.94** | **0.022\*** |
|  | Chemotype 1 – Chemotype 3 | **9.40** | **0.007\*\*** |
|  | Chemotype 2 – Chemotype 3 | 0.38 | 0.539 |
| Total root length | Chemotype 1 – Chemotype 2 | **4.94** | **0.039\*** |
|  | Chemotype 1 – Chemotype 3 | **13.01** | **0.001\*\*** |
|  | Chemotype 2 – Chemotype 3 | 1.83 | 0.176 |
| Volume | Chemotype 1 – Chemotype 2 | **6.80** | **0.027\*** |
|  | Chemotype 1 – Chemotype 3 | 4.35 | 0.055. |
|  | Chemotype 2 – Chemotype 3 | 0.26 | 0.611 |
| Root:shoot ratio | Chemotype 1 – Chemotype 2 | **11.49** | **0.001\*\*** |
|  | Chemotype 1 – Chemotype 3 | **16.92** | **<0.001\*\*\*** |
|  | Chemotype 2 – Chemotype 3 | 0.50 | 0.479 |

